# Preclinical Characterization of Relatlimab, a Human LAG-3–Blocking Antibody, Alone or in Combination With Nivolumab

**DOI:** 10.1101/2022.01.24.477551

**Authors:** Kent Thudium, Mark Selby, Julie A. Zorn, Gregory Rak, Xi-Tao Wang, Roderick Todd Bunch, Jason M. Hogan, Pavel Strop, Alan J. Korman

## Abstract

Novel therapeutic approaches combining immune checkpoint inhibitors are needed to improve clinical outcomes for patients with cancer. Lymphocyte-activation gene 3 (LAG-3) is an immune checkpoint molecule that inhibits T-cell activity and antitumor immune responses, acting through an independent mechanism from that of programmed death-1 (PD-1) and cytotoxic T lymphocyte associated antigen-4 (CTLA-4). Here, we describe the development and preclinical characterization of relatlimab, a human antibody that binds to human LAG-3 with high affinity and specificity to block the interaction of LAG-3 with the ligands MHC II and fibrinogen like protein-1, and to reverse LAG-3–mediated inhibition of T-cell function *in vitro*. Consistent with previous reports, in mouse models, the combined blockade of LAG-3 and PD-1 with surrogate antibodies resulted in enhanced anti-tumor activity greater than the individual blockade of either receptor. In toxicity studies in cynomolgus monkeys, relatlimab was generally well tolerated when combined with nivolumab. These results are consistent with findings from the RELATIVITY-047 phase 2/3 trial showing that relatlimab combined with nivolumab is a well-tolerated regimen that demonstrated superior progression-free survival compared with nivolumab monotherapy in patients with unresectable or metastatic melanoma.

**Synopsis:** Preclinical studies demonstrate that relatlimab specifically blocks the interaction between LAG-3 and its ligands, and provide a biological rationale for combining relatlimab with the anti−PD-1 antibody nivolumab as an effective cancer immunotherapeutic strategy.

## Introduction

Immune checkpoint blockade (ICB) has revolutionized treatment options for patients with cancer, improving the survival of patients with a range of different malignancies (1). An effective immune response against cancer relies on immune surveillance of tumor antigens expressed on cancer cells, which ultimately results in an adaptive immune response and cancer cell death (2-4). Tumor immune escape, facilitated in part by the expression of inhibitory ligands, can be reversed by the blockade of receptors such as programmed death-1 (PD-1) and cytotoxic T lymphocyte antigen-4 (CTLA-4), but there remains a need for additional novel combinations to improve patient outcomes (5-7).

Lymphocyte-activation gene 3 (LAG-3) is an inhibitory receptor on T cells frequently upregulated on the surface of tumor infiltrating lymphocytes (TILs), including regulatory T cells, which contributes to T-cell exhaustion in the tumor microenvironment (TME) and limits antitumor T-cell responses (8-11). The mechanism by which LAG-3 exerts its immune suppressive effects is not completely understood but includes engagement by MHC II expressed on antigen-presenting cells (APCs) within the TME and on some tumor cells, as well as potential engagement by other reported ligands, including fibrinogen like protein-1 (FGL1) (5,12).

Previous studies have demonstrated that LAG-3 and PD-1 act in a nonredundant manner to suppress T-cell stimulation. In an *in vitro* antigen-specific T-cell stimulation system, the peptide responsiveness of T cells transduced to express both LAG-3 and programmed death ligand 1 (PD-L1) showed lower levels of interleukin-2 (IL-2) secretion in co-culture with APCs expressing PD-L1 and MHC II compared with the responses of T cells expressing either receptor alone (13). Preclinical data has demonstrated a synergistic relationship between the inhibitory receptors LAG-3 and PD-1 in regulating immune homeostasis, preventing autoimmunity, and enforcing tumor-induced tolerance (9,13). Importantly, in mice, antibody blockade of both receptors resulted in more robust immune responses compared with blockade of either individual receptor (14-16).

Here, we describe the development of relatlimab, a human LAG-3 (hLAG-3) blocking antibody, and preclinical analyses that demonstrate the binding affinity, specificity, functional activity, and safety of this novel immune checkpoint inhibitor. *In vitro* assay results were consistent with published *in vivo* data showing that LAG-3 blockade combined synergistically with PD-1 blockade to achieve enhanced antitumor and immunomodulatory activity. We observed a weak but measurable interaction *in vitro* between FGL1 and LAG-3, and confirmed both the inhibitory potential of this interaction as well as the ability of relatlimab to block it. Our overall findings are consistent with the results from the RELATIVITY-047 study, the first phase II/III trial evaluating dual administration of relatlimab and nivolumab in patients with previously untreated or unresectable melanoma. In this trial, the combined blockade of LAG-3 and PD-1 demonstrated superior progression-free survival (PFS) compared with blockade of PD-1 alone (17,18).

## Materials and Methods

### Relatlimab generation and characterization

#### Generation of human monoclonal antibody anti–LAG-3 antibody (relatlimab)

Transgenic mice comprising germline configuration human immunoglobulin (Ig) miniloci in an endogenous IgH and Igκ knockout background (19,20) were immunized with recombinant human LAG-3–Fc protein (hLAG-3–hFc) (R&D Systems, Minneapolis, MN, USA), consisting of the extracellular domain of LAG-3 (Leu23-Leu450) fused to the Fc portion of human IgG1, together with Ribi adjuvant (Ribi lmmunoChemical Research, Hamilton, MT, USA). Spleen cells from immunized mice were fused with P3×63Ag8.653 myeloma cells (ATCC, Manassas, VA, USA) and screened for hybridomas-producing human monoclonal antibodies (mAbs) reactive to LAG-3–Fc by enzyme-linked immunosorbent assay (ELISA). Purified antibodies against hLAG-3, including clone 25F7, were analyzed by flow cytometry for the potency of binding to transfected Chinese hamster ovary (CHO) suspension cells overexpressing hLAG-3 (Bristol Myers Squibb, Princeton, NJ, USA), but not to parental CHO cells. The variable region sequences of clone 25F7 were cloned and subsequently grafted onto human κ and IgG4 constant region sequences for recombinant CHO cell expression of the resulting derived antibodies, LAG3.1-G4P and LAG3.5-G4P (relatlimab) (see Results for additional detail).

### Epitope characterization

The relatlimab-binding epitope on LAG-3 was determined by several methods including peptide library binding, binding to LAG-3 truncation mutants, differential chemical labeling, and x-ray crystallography. Detailed descriptions of the methods employed are presented in the Supplementary Methods.

### Octet^®^ biolayer interferometry characterization of relatlimab blockade of LAG-3/MHC II and LAG-3/FGL1 interactions

#### LAG-3/MHC II interaction and blockade by relatlimab

To assess the blockade of human leukocyte antigen DR1 isotype (HLA-DR1)/hLAG-3–hFc engagement by either divalent relatlimab or its antigen-binding fragment (Fab) by Octet^®^ biolayer interferometry (BLI), hLAG-3–hFc was first incubated with excess relatlimab or Fab 3.5 in bovine serum albumin in phosphate buffered saline-tween (PBST-BSA). Then, biotinylated HLA-DR1 was captured on Streptavidin biosensors (ForteBio, CA, USA) at 10 μg/mL for 180 seconds in PBST-BSA followed by a 60-second wash with PBST-BSA. hLAG-3–hFc:relatlimab or hLAG-3–hFc:Fab 3.5 complexes were tested for binding to the captured HLA-DR1 protein over 300 seconds.

#### LAG-3/FGL1 interaction and blockade by relatlimab

The binding of recombinant human FGL1, fused with mouse Fc at the N-terminus (mFc-FGL1 and mFc-FGL1-FD) to recombinant hLAG-3–hFc was investigated using BLI. Detailed methods are provided in the Supplementary Methods.

### Immunohistochemistry analysis in normal human tissues

Immunohistochemistry (IHC) analyses were performed in a selected panel of normal human tissues to detect tissue binding. Detailed methods are provided in the Supplementary Methods.

### Toxicity studies in cynomolgus monkey model

#### Study design

As part of a 4-week multi-dose toxicity study, monkeys were dosed once weekly (QW) for five total doses with relatlimab alone or in combination with nivolumab. Serum concentrations of relatlimab and nivolumab were determined from blood samples collected from all available monkeys at multiple timepoints between days 1 and 71 of the study. The clinical evaluation parameters and dosing strategy for this study are described in Supplementary Table S1.

The potential toxicity of relatlimab was also assessed in a 3-month multi-dose toxicity study in monkeys administered drug QW for 3 months. Relatlimab was administered by intravenous infusion at doses of 0 (vehicle), 30, and 100 mg/kg to groups of six monkeys/sex. The clinical evaluation parameters and dosing strategy for this study are described in Supplementary Table S2.

### Statistical analyses

Statistical analyses of the peripheral blood lymphocyte phenotyping, keyhole limpet hemocyanin (KLH)-specific antibodies, and splenic T-lymphocyte subset phenotyping data were performed by Global Biometric Sciences Nonclinical Biostatistics. Appropriate nonstatistical methods were used for group comparisons of immunogenicity data and *ex vivo* recall responses to KLH and hepatitis B surface antigen.

### Additional methods

Additional details on methods and experiments relating to relatlimab binding to human and cynomolgus monkey LAG-3, binding affinity, cell-based bio- and blocking assays, antibody-dependent cellular cytotoxicity (ADCC), polymerase chain reaction (PCR) analysis, splenic T-lymphocyte subset phenotyping, *ex vivo* responses to KLH-specific antibodies, study animals, antibodies used, cell cultures, and procurement of human tissue samples can be found in the Supplementary Methods.

## Data Availability

Selected datasets in support of the findings of this study will be deposited to suitable repositories and made available to readers of *Cancer Immunology Research* at manuscript acceptance and ahead of publication. X-ray diffraction coordinates and structure factors will be deposited in the Research Collaboratory for Structural Bioinformatics Protein Data Bank under accession code XXXX. (Note to editor and peer-reviewers: Data will be made available to the editor and peer-reviewers upon request).

## Results

### Relatlimab antibody generation

A panel of human anti–LAG-3 mAbs was generated by immunization of HuMAb^®^ and KM^®^ transgenic mice with recombinant LAG-3 protein. Hybridoma clone 25F7 was selected for expansion and further characterization based on its activity profile in biochemical and cell-based binding and blocking assays, as well as in functional T-cell activation assays. The variable region sequences of this antibody were cloned and subsequently grafted onto human κ and IgG4 constant region sequences for reduced Fc receptor engagement and a reduced potential for effector T-cell mediated cytotoxicity. The resulting antibody, LAG3.1-G4P, also incorporated a S228P stabilizing hinge mutation to prevent *in vivo* and *in vitro* IgG4 Fab-arm exchange (21).

The sequence of antibody LAG-3.1-G4P contained two potential deamidation sites in the CDR2 region of the heavy chain at residues N54 and N56, which were confirmed by biophysical analysis under forced deamidation conditions. These potential sequence liabilities were addressed by mutating these sites (N54R and N56S), with no deleterious effect observed on the functional activity of the resulting antibody designated LAG-3.5, later renamed BMS-986016 (relatlimab) (data not shown).

### Relatlimab-binding specificity

Relatlimab displayed saturable and selective binding to immobilized hLAG-3–hFc by ELISA, with a half-maximal effective concentration (EC_50_) of 0.49 nM compared with an EC_50_ of 1.46 nM for binding to CHO cells expressing hLAG-3 (Supplementary Fig. S1A). Relatlimab did not bind to recombinant mouse LAG-3 protein or to cells expressing full-length murine LAG-3 (data not shown). To confirm that relatlimab recognized native LAG-3, the binding of relatlimab to primary activated human and cynomolgus CD4+ T cells was measured and found to be substantially higher for binding to human (mean EC_50_, 0.11 nM) than monkey LAG-3 (mean EC_50_, 29.11 nM) (**Fig. 1A** and **B**).

**Figure 1.**
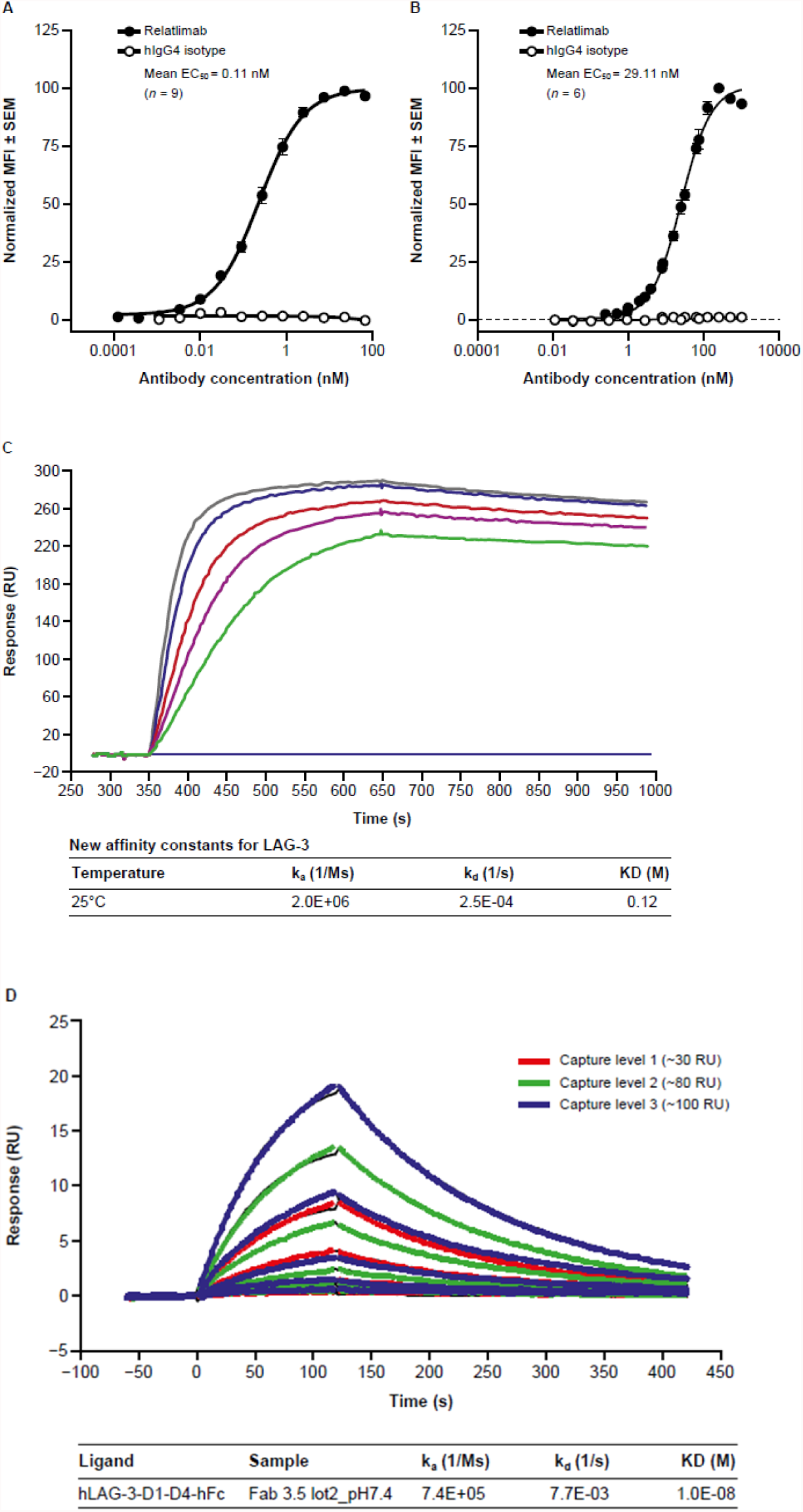
Characterization of LAG-3 binding by relatlimab. **A**, Normalized binding of relatlimab to activated human CD4+ T cells from nine donor samples. **B**, Normalized binding of relatlimab to activated cynomolgus CD4+ T cells from six donor samples. **C**, Sensogram depicting binding of bivalent relatlimab to LAG-3 under neutral pH conditions. **D**, Sensogram depicting binding of monomeric relatlimab Fab under neutral pH conditions. hIgG4, human IgG4; KD, dissociation constant; MFI, mean fluorescence intensity.

The kinetics of relatlimab binding to LAG-3 were determined by surface plasmon resonance (SPR) for both intact bivalent relatlimab and its Fab fragment. The apparent affinity of the bivalent antibody was found to be 0.12 nM at pH 7.4 under the experimental conditions detailed in the Supplementary Methods. The monovalent affinity of the Fab fragment measured with SPR was 10 nM (**Fig. 1C** and **1D**). Binding at pH 6, acidic conditions that likely exist in the TME (22), shows faster dissociation kinetics with a modest effect on affinity, indicating that relatlimab would still bind under these conditions (Supplementary Fig. S1B).

### Disruption of LAG-3 receptor/ligand interactions by relatlimab

The functional potency of relatlimab in blocking the interaction of LAG-3 with MHC II and FGL1 was examined in a series of biochemical and cell-based assays.

#### MHC II binding

The interaction of LAG-3 with MHC II was confirmed by comparing the binding of mouse and human LAG-3-hFc to the wild-type Raji B lymphoid cell line and to a MHC II-low variant, RJ225 (23), with substantially less binding observed to the mutant cell line compared with the wild-type line (Supplementary Fig. S2). Relatlimab completely blocked detectable binding of LAG-3–mFc to MHC II+ Daudi B lymphoid cells, exhibiting a half-maximal blockade (IC_50_) of 0.67 nM compared with isotype control antibody (**Fig. 2**). Similar results were also obtained using wild-type Raji B lymphoid cells (data not shown). Blockade was confirmed by BLI measurement of the binding of LAG-3-Fc to HLA-DR1 in the presence of either bivalent relatlimab or a Fab fragment of relatlimab. The interaction was equivalently blocked by both relatlimab and its Fab, but not by other antibody clones that bind within the D3-D4 extracellular domains (Supplementary Fig. S3).

**Figure 2.**
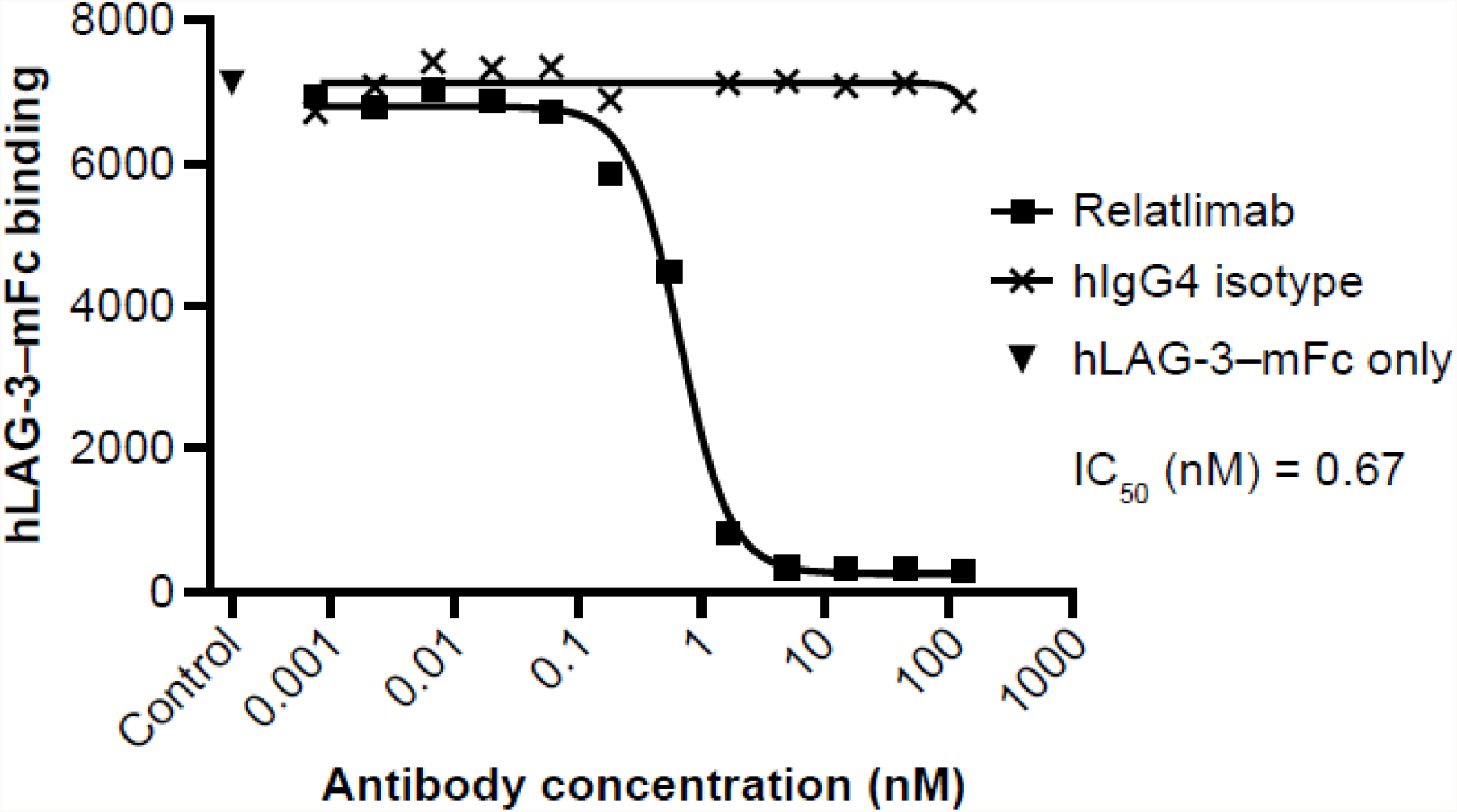
Characterization of *in vitro* interactions of relatlimab. Blockade by relatlimab of LAG-3–mFc binding to MHC II+ Daudi cells.

#### FGL1 binding

We next evaluated whether relatlimab can block the interaction between LAG-3 and the recently identified ligand, FGL1, using ELISA. Recombinant hLAG-3–hFc fusion protein bound immobilized recombinant human FGL1-mFc (hFGL1-mFc) protein (3 µg/mL), with an EC_50_ of 0.085 nM (**Fig. 3A**) and was blocked by relatlimab, with an IC_50_ of 0.019 nM, but not by the isotype control antibody (**Fig. 3A** and **B**).

**Figure 3.**
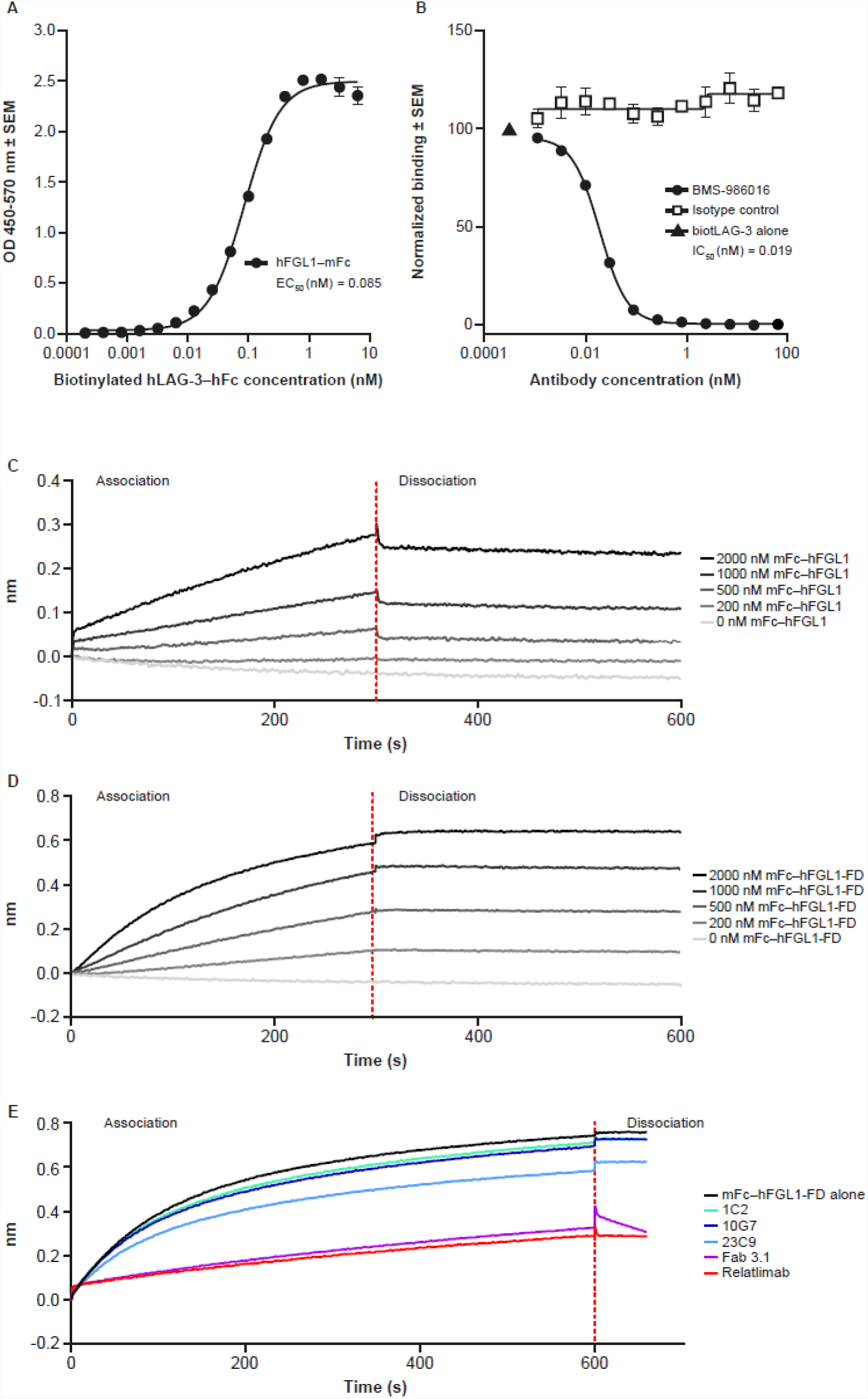
Measurement of LAG-3/FGL1 interaction and blockade. **A**, Binding of hLAG-3 to mFc–hFGL1. **B**, Blockade of hFGL1/LAG-3 engagement by relatlimab. **C**, Binding of mFc– hFGL1 to hLAG-3. **D**, Binding of mFc–hFGL1-FD (fibrinogen domain alone) to hLAG-3. **E**, Blockade of mFc–hFGL1-FD (fibrinogen domain alone)/LAG-3 engagement by relatlimab.

Similar studies were conducted using BLI, which confirmed that both full-length FGL1 (mFc–hFGL1), as well as the fibrinogen domain alone (mFc–hFGL1-FD), bound hLAG-3-hFc in a dose-dependent, albeit nonsaturable, manner up to 2000 nM FGL1, indicating a weak interaction (**Fig. 3C** and **D**). This interaction was inhibited by relatlimab (**Fig. 3E)**. While the mechanism by which soluble FGL1 may productively engage LAG-3 to drive T-cell inhibition is unknown, these results corroborate those of Wang et al., suggesting that the FGL1 fibrinogen domain alone is sufficient to bind LAG-3 (12).

### Functional blockade of ligand interaction with LAG-3

The ability of human LAG-3 to inhibit T-cell responses was studied using an antigen-specific T-cell hybridoma, 3A9, specific for hen egg lysozyme peptide (HEL48-62), presented by the MHC II-matched antigen-presenting mouse cell line LK35.2 (24). The expression of full-length hLAG-3 on the T cells (3A9-hLAG-3) resulted in attenuated T-cell peptide-responsiveness, as demonstrated by lower murine IL-2 secretion when co-cultured with LK35 cells that could be enhanced by relatlimab in a dose-dependent manner (**Fig. 4A**).

**Figure 4.**
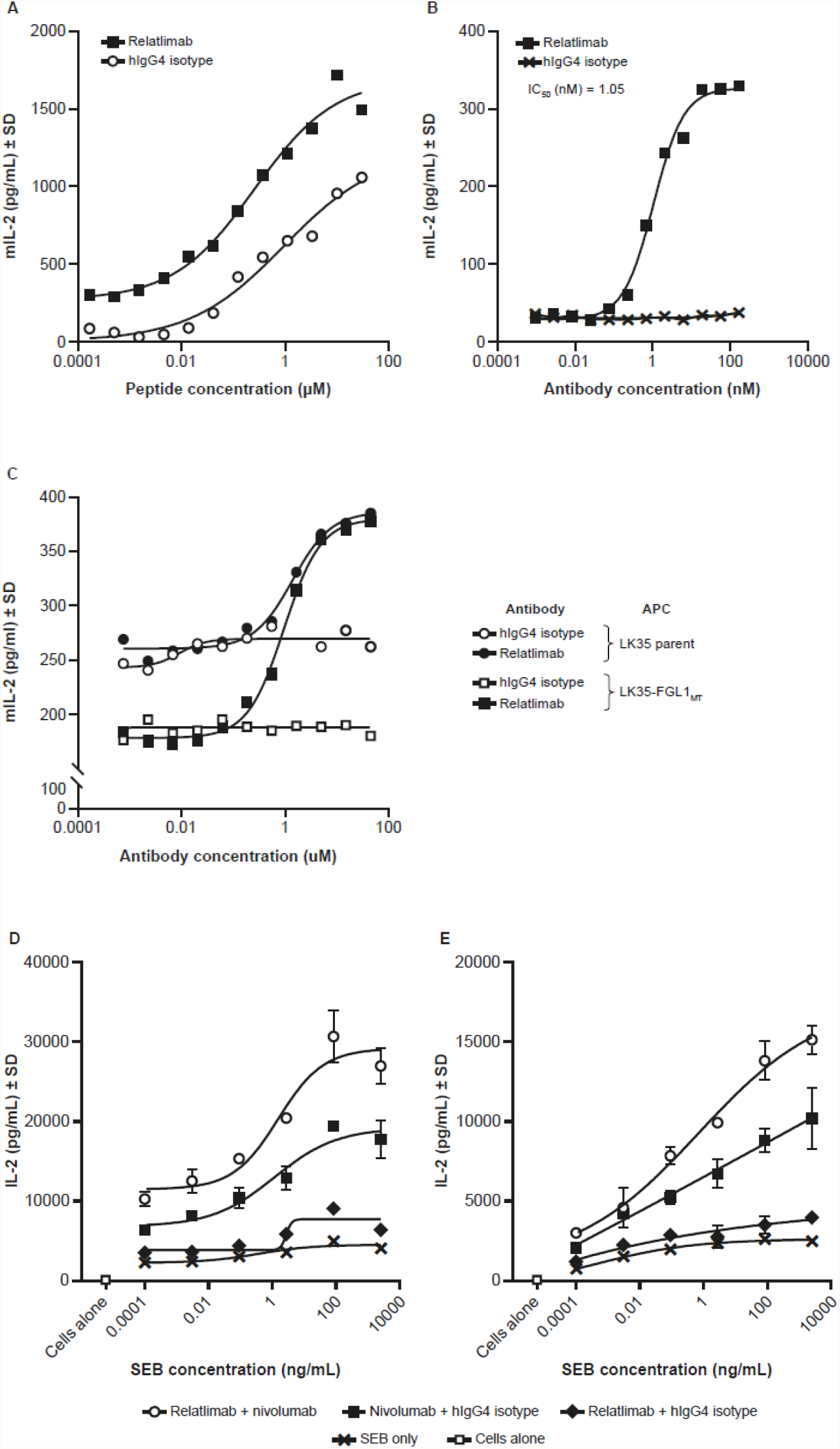
Characterization of *in vitro* interactions of relatlimab. **A**, Enhanced peptide responsiveness of T-cell hybridoma 3A9-hLAG-3 by relatlimab. **B**, Reversal of LAG-3–mediated inhibition of T-cell hybridoma 3A9-hLAG-3 by relatlimab. **C**, Measurement of LAG-3/FGL1 interaction and blockade by relatlimab of FGL1- and MHC II-mediated T-cell inhibition in the 3A9-hLAG-3 T-cell hybridoma assay. **D** and **E**, Enhanced activation of superantigen-stimulated human PBMC cultures from donor 1 (**D**) and donor 2 (**E**) by relatlimab alone or in combination with nivolumab with examples from two donors. mIL-2, mouse interleukin 2; MT, membrane tethered.

Using an alternate format of the assay where cells were stimulated with a suboptimal concentration of HEL48-62 peptide in the presence of titrated mAb, relatlimab displayed potent blockade of LAG-3-mediated inhibition (IC_50_, 1.05 nM) compared with the isotype control antibody (**Fig. 4B**). Collectively, these results suggest that the observed functional T-cell inhibition seen in co-culture with LK35 cells and antigen, which is likely mediated by murine class II MHC II present on the APCs interacting with hLAG-3 on the T cells, can be reversed by relatlimab in a dose-dependent manner.

In contrast to published results (12), we observed only modest T-cell inhibitory activity by soluble mFc–hFGL1 that had been purified to ensure a homogenous monomeric preparation (Supplementary Fig. S4) and, consequently, we next evaluated whether higher order oligomers of FGL1 could elicit enhanced inhibition. We generated an ordered oligomer of FGL1 by fusion to hexamer-forming Fc (E345R/E430G/S440Y) (FGL1-RGY) (25,26). While increased T-cell inhibition was observed with this variant relative to the wild-type Fc fusion protein alone, the impact on activity was still modest (Supplementary Fig. S4). For this reason, we sought to determine whether expression of a membrane-tethered version of FGL1 on the cell surface could allow for a higher avidity interaction of FGL1 with LAG-3, thereby revealing functional inhibitory engagement. A membrane-tethered version of FGL1 (FGL1_MT_) was generated by fusing the fibrinogen domain of FGL1 with the transmembrane and intracellular domain of type II membrane protein FIBCD (12), and was expressed on LK35 cells (LK35-FGL1_MT_) (Supplementary Fig. S5). 3A9-hLAG-3 cells were more attenuated in their peptide responsiveness in co-culture with LK35-FGL1_MT_ cells compared with parental LK35 cells expressing MHC II alone (**Fig. 4C**). This suggests that FGL1-mediated inhibition in this setting is additive to the inhibition resulting from MHC II/LAG-3 engagement. Relatlimab reversed the T-cell inhibition resulting from co-culture with either parental LK35 and LK35-FGL1 cells with similar potency (IC_50_ of 1.39 nM and 0.95 nM, respectively) and to equivalent maximal levels of cytokine production, suggesting that relatlimab can effectively block the simultaneous engagement of LAG-3 by both MHC II and FGL1 (**Fig. 4C** and Supplementary Fig. S5).

The functional activity of relatlimab in the context of primary T cells was evaluated in peripheral blood mononuclear cell (PBMC) cultures stimulated with superantigen Staphylococcal enterotoxin B. Cultures from 15 of 18 donors showed enhanced IL-2 secretion in the presence of relatlimab alone compared with the isotype control and, in most instances, the stimulation was less than that observed for treatment with nivolumab. The combination of relatlimab and nivolumab resulted in higher levels of stimulation compared with a combination of nivolumab and isotype control (**Fig. 4D** and **E**).

Finally, relatlimab binding to activated T cells did not mediate significant levels of ADCC when compared with the isotype control, whereas binding with positive control nonfucosylated human IgG1 anti-CD30 antibody resulted in robust cell lysis (Supplementary Fig. S6).

These results are consistent with the synergistic *in vivo* antitumor activity observed from the combination antibody blockade of mouse LAG-3 and PD-1 in syngeneic tumor models. Similar to previously published reports (9), the antitumor activity of anti–mouse-LAG-3 mAbs C9B7W and 19C7, in combination with anti–mouse-PD-1 monoclonal antibody 4H2, showed enhanced efficacy when compared with LAG-3 or PD-1 single-agent blockade in both MC38 colon carcinoma tumors (Supplementary Fig. S7) and in Sa1N fibrosarcoma tumors (Supplementary Fig S8). Substantial antitumor activity from the blockade of LAG-3 alone was only observed in the immunogenic Sa1N model.

### Characterization of the relatlimab binding site

Previous studies by Triebel et al. (27) demonstrated that an extra insertion-loop sequence in the N-terminal extracellular D1 of LAG-3 potentially mediates the interaction with MHC II (28). Our initial analysis of hybridoma clones revealed that the most potent antibodies, including the parent clone for relatlimab (LAG3.1-G4P), were bound within the N-terminal D1-D2 domain region of LAG-3 (Supplementary Fig. S3C). Interestingly, when a peptide consisting of the full-length D1 insertion-loop sequence (P_60_GPHPAAPSSWGPRPR_75_) was synthesized and tested for antibody reactivity, LAG3.1-G4P was observed to bind strongly to this peptide by ELISA, with an EC_50_ of 0.44 nM (data not shown). Next, LAG3.1-G4P was assessed by ELISA for binding a set of overlapping peptides spanning the 30 residues of the LAG-3 insertion loop (Supplementary Fig. 9A) and indicated that the antibody binds residues within the peptide H_63_PAAPSSW_70_ (**Fig. 5A**). Carbene chemical footprinting of a complex consisting of the LAG-3 D1-D2 domains and relatlimab Fab confirmed that the epitope is contained within the peptide A59-W70, since this peptide showed the largest decrease in chemical labeling following complex formation compared with LAG-3 D1-D2 alone (**Fig. 5B, upper panel**). Residue level labeling of peptide A59-W70 identified multiple amino acids undergoing protection following complex formation contained in the region H63-W70 (**Fig. 5B, lower panel**). Additional peptides were observed with labeling protection that was localized to individual amino acids in the D1 domain. As the protein structure of LAG-3 with the insertion loop has not been determined, we speculate that these residues are likely to be structurally located near the A59-W70 peptide. Comparison of the insertion-loop sequences of human and cynomolgus monkey LAG-3 proteins showed 75% sequence identity for the epitope (H63–W70), likely accounting for the lower affinity of relatlimab binding to nonhuman primate LAG-3 (Supplementary Fig. 9B).

**Figure 5.**
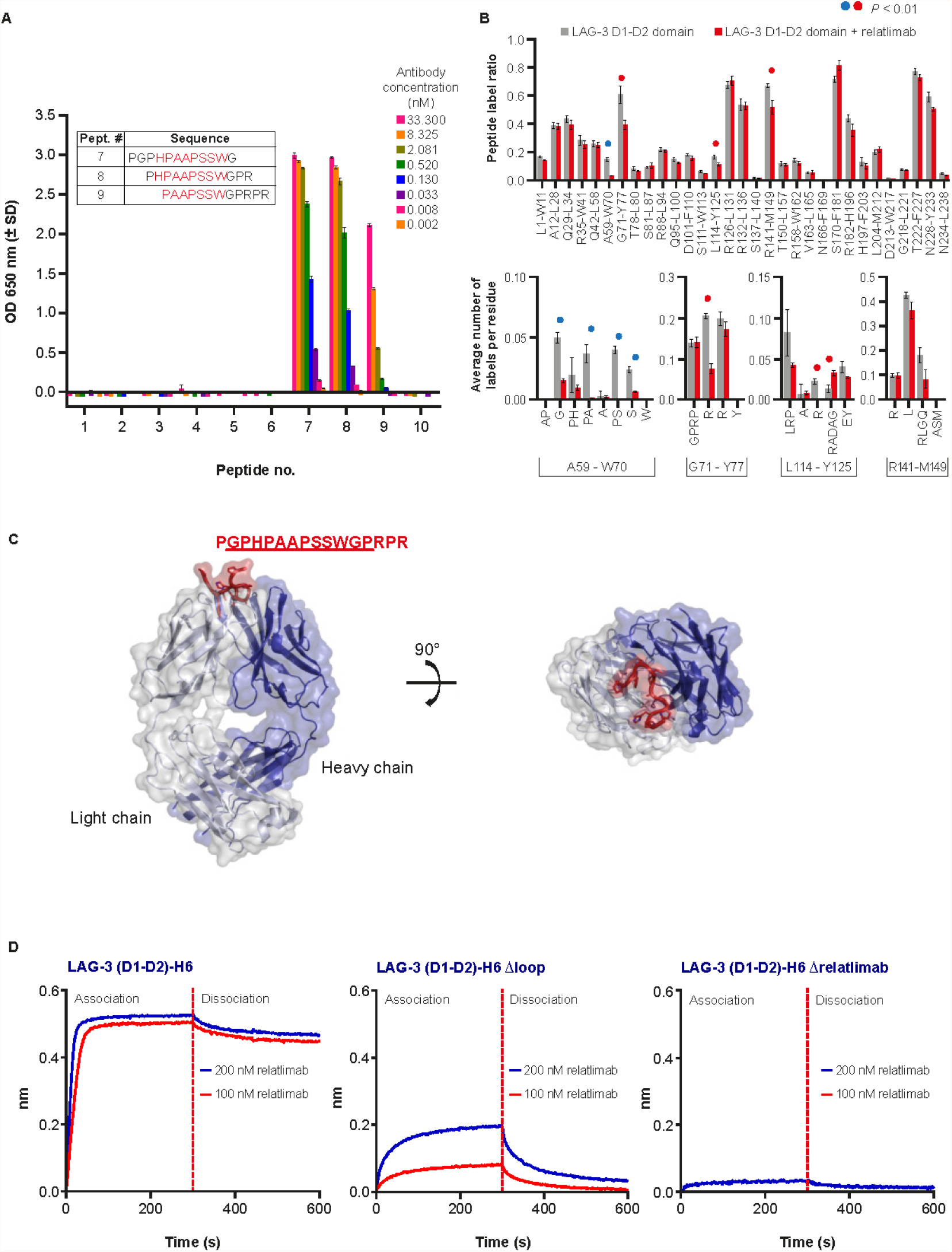
Relatlimab epitope characterization. **A**, Epitope determination by overlapping peptide ELISA. **B**, Mapping of relatlimab epitope by carbene chemical footprinting. Blue circles denote the binding site peptide A59-W70. Red circles denote nearby residues in the D1 domain with some protection but not the physical epitope. **C**, X-ray structure of the LAG3.1-G4P Fab complex with binding peptide epitope. Density is observed for the underlined sequence. **D**, Binding of human LAG-3 loop deletion variants to relatlimab by BLI.

### Structural characterization of relatlimab in complex with the LAG-3 insertion loop

To further characterize relatlimab interactions with LAG-3, the structure of the Fab of LAG3.1-G4P in complex with a 16-residue peptide derived from insertion-loop sequence of the LAG-3 D1 domain was determined using x-ray crystallography (**Supplementary Table S3**). The three-dimensional structure, captured at 2.4 Å resolution, revealed distinct density for 12 of the 16 peptide residues (called out sequence in **Fig. 5C**) in the LAG3.1-G4P binding groove, with CDR-H3 and CDR-L3 pushed apart (**Fig. 5C**). Half of the accessible surface area of the peptide is buried by LAG3.1-G4P, with light chain interactions more prevalent over heavy chain interactions (380 Å2 vs. 160 Å2, respectively). Roughly half (680 Å2) of the total accessible surface area of the peptide (1350 Å2) is buried by LAG3.1-G4P. Protein-binding assays using LAG-3 domain variants further confirmed this core peptide sequence as essential for the interaction between relatlimab and LAG-3, with deletion of the H63–W70 sequence resulting in the abolishment of relatlimab binding (**Fig. 5D** and Supplementary Fig. S9C). Overall, the peptide binding, BLI, and chemical footprinting data are in good agreement and are corroborated by the x-ray crystal structure data and LAG-3 domain variant-binding data to collectively identify a linear epitope of relatlimab centered on residues H63–W70 of the insertion loop.

### Relatlimab tissue-binding properties in normal human tissues

To confirm LAG-3 expression in immune cells, and to assess any unexpected tissue binding, tissue cross-reactivity of relatlimab and the parent clone of relatlimab, LAG3.1-G4P, was determined in a selected panel of normal human tissues by IHC (Supplementary Table S4). The tissue-binding patterns by relatlimab and LAG3.1-G4P were very similar. In hyperplastic tonsil tissue, strong positive staining was revealed in a small subset of lymphocytes primarily distributed in the interfollicular area (T-cell region), with few in the mantle zone, and only extremely rarely in the germinal center (**Fig. 6A**). This lymphocyte staining was expected and consistent with published observations (29). In pituitary tissue, rare to occasional moderate/strong immunoreactivity was displayed in the adenohypophysis. No specific staining was observed in the other tissues examined (Supplementary Table S4). Dual-color immunofluorescence analysis with LAG3.1-G4P confirmed the specificity of this staining localization of LAG-3 in pituitary gonadotroph cells, which include follicular-stimulating hormone- and luteinizing hormone (LH)-producing cells. Pituitary expression of LAG-3 was also confirmed by PCR (**Fig. 6A** and **B**; Supplementary Fig. S10 and S11, Supplementary Table S4).

**Figure 6.**
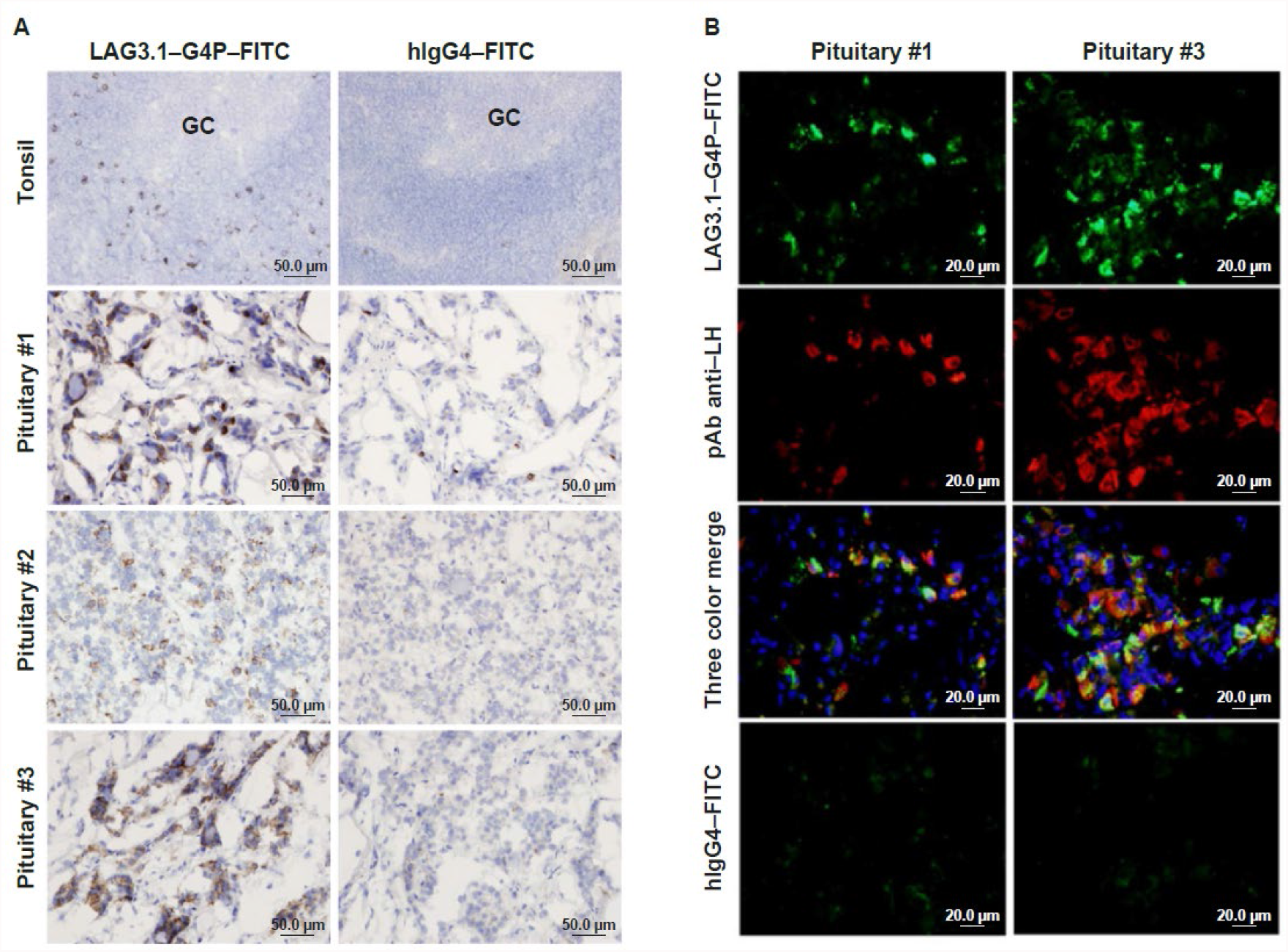
Binding of LAG3.1-G4P in human pituitary gland. **A**, IHC of FITC-conjugated LAG-3.1-G4P (left panels) in positive control tissue (hyperplasia tonsil) and three normal human pituitary samples. FITC-conjugated human IgG4 was used as isotype control (right panels). GC, germinal center of the tonsil. Counterstained with Mayer’s hematoxylin. **B**, Co-localized binding of HuMAb^®^ anti–LAG-3 in LH-producing gonadotroph cells. Double immunofluorescent staining of FITC-conjugated LAG-3.1-G4P (green) and rabbit pAb anti-LH (red) on cryostat sections from two normal human pituitary samples. FITC-conjugated human IgG4 and rabbit IgG (not shown) were used as isotype controls (bottom panels) while nuclei were counterstained using Hoechst 33342. FITC, fluorescein isothiocyanate; pAb, polyclone antibody.

### Preclinical toxicity assessment in cynomolgus monkeys

As relatlimab was shown to bind to cynomolgus LAG-3, we evaluated the potential toxicity of relatlimab ± nivolumab in cynomolgus monkeys. Additionally, splenic T-cell phenotyping, T-cell phenotyping, and T-cell dependent antibody responses (TDARs) were analyzed (Supplementary Methods).

#### Repeat-dose toxicity

In a 4-week repeat-dose toxicity study (Supplementary Table S1), relatlimab was clinically well tolerated by cynomolgus monkeys when administered intravenously QW at 30 or 100 mg/kg with no adverse findings. Relatlimab, when administered at 100 mg/kg in combination with nivolumab at 50 mg/kg, was generally well tolerated in eight out of nine monkeys with no adverse clinical signs, the exception being moribundity in one male monkey attributed to central nervous system (CNS) vasculitis (Supplementary Fig. S12). Additional minimal histopathological findings in the combination group were likely the result of enhanced immunostimulatory effects of nivolumab in combination with relatlimab, since no treatment-related histopathological changes were noted with relatlimab alone (Supplementary Table S5). The CNS vascular findings may have been a result of a loss of tolerance to self-antigens based on the synergistic role of PD-1 and LAG-3 in maintaining self-tolerance. Given the long half-lives of relatlimab and nivolumab, the irreversibility of relatlimab plus nivolumab-related findings is likely a result of continued exposure to the test articles throughout the duration of the recovery period. In a 3-month toxicity study (Supplementary Table S2), relatlimab was generally well tolerated by mature cynomolgus monkeys (4 to 7 years old) when administered intravenously QW up to 100 mg/kg.

### In vivo *pharmacodynamic effects of relatlimab administration*

As part of the 4-week repeat-dose toxicity study of relatlimab and nivolumab in monkeys, all animals were immunized intramuscularly on day 1 of the study with 10 mg of KLH to permit assessment of *in vivo* antibody responses, *ex vivo* recall responses to KLH, and immunophenotypic analyses of peripheral blood and splenic T-lymphocyte subsets.

There were no observed relatlimab- or nivolumab-related changes in TDARs to KLH among any of the study groups, and no *ex vivo* recall responses to KLH in CD8+CD4-T cells. The comparatively low affinity of relatlimab for cynomolgus LAG-3 may have resulted in incomplete receptor blockade resulting in attenuated lymphocyte responses, including TDAR. Nevertheless, drug-related changes in *ex vivo* T-cell recall responses to KLH observed *in vitro* were indicative of an enhanced antigen-specific response. These included increases at day 22, which waned by day 57, in the mean percentage of (1) CD69^+^, TNF-α^+^, and CD69^+^TNF-α^+^ CD4^+^CD8^-^ splenic T cells in female monkeys in all dose groups and in males with 100 mg/kg relatlimab alone or in combination with nivolumab without meaningful differences in degree between the two regimens (**Fig. 7A** and **B)**; and of (2) CD69^+^TNF-α^+^IFN-γ^+^ CD4^+^CD8^-^ splenic T cells in both sexes at 100 mg/kg relatlimab and 50 mg/kg nivolumab when administered individually, with further non-additive increases in the combination group (Supplementary Fig. S13). Changes in lymphocyte phenotypes at 100 mg/kg relatlimab alone compared with control monkeys were limited to increases in mean percentage CD25^+^FoxP3^+^ CD8^+^ T cells. An increase in the mean percentage of CD25^+^FoxP3^+^ CD8^+^ T cells (1.6 to 1.9× relative to vehicle control) was found on day 30 in monkeys administered with nivolumab 50 mg/kg/week ± relatlimab 100 mg/kg/week, which continued into the recovery period on day 72 (1.5 to 1.6× relative to control) (**Fig. 7C**).

**Figure 7.**
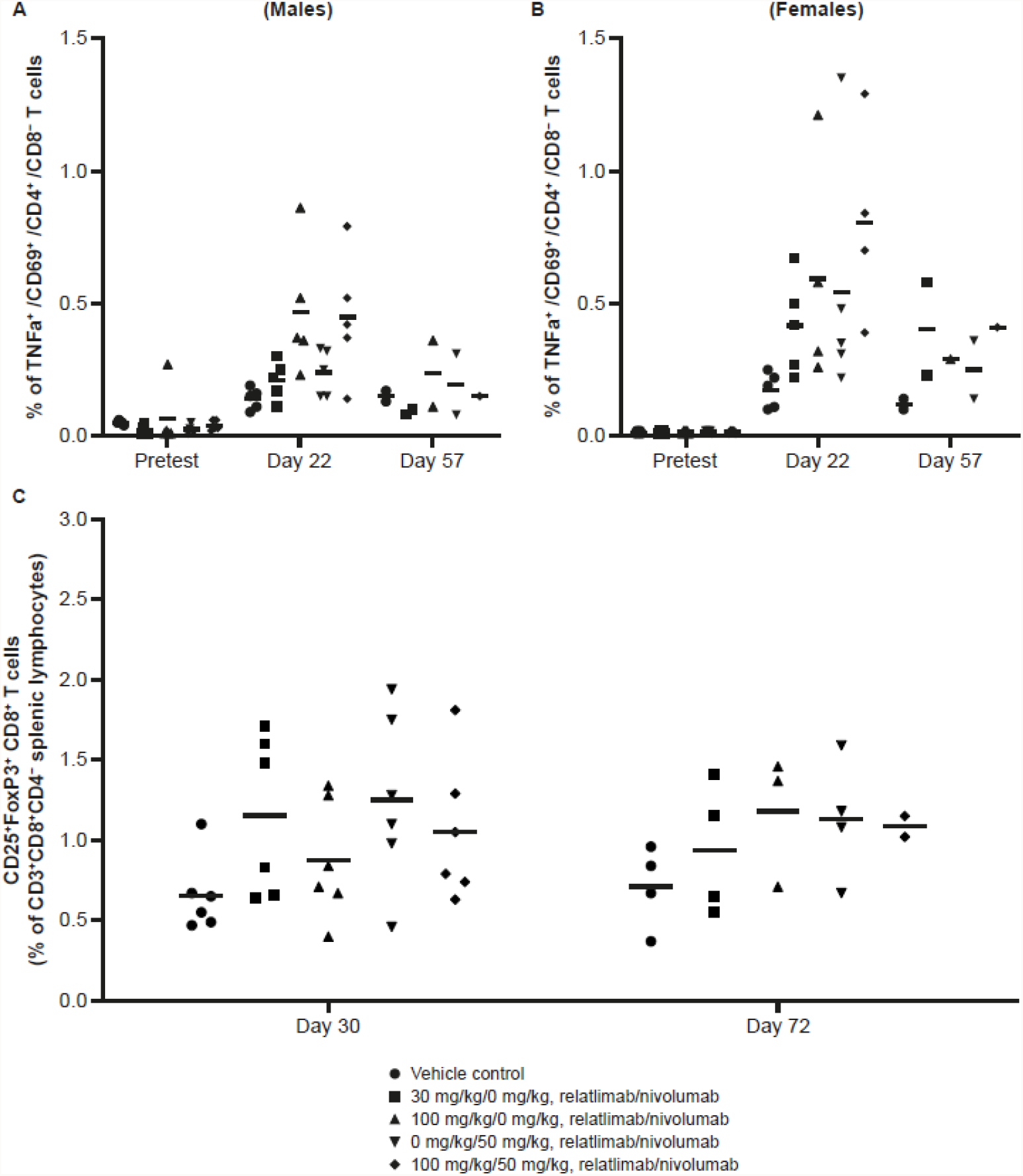
Changes to splenic T-lymphocytes and CD69^+^TNF-α^+^ CD4^+^CD8-T-cell populations in monkeys treated with relatlimab ± nivolumab. **A** and **B**, *Ex vivo* recall response to KLH results for CD69^+^TNF-α^+^CD4^+^CD8^-^ T cells in male (**A**) and female (**B**) monkeys. Data points represent individual animal results as a percentage of CD4^+^CD8^-^ cells. Horizontal bars represent group means. **C**. Splenic T-lymphocyte subset phenotyping results for CD25^+^FoxP3^+^CD8 T cells. Data points represent individual animal results (males and females) as a percentage of CD3^+^CD8^+^CD4-splenic lymphocytes. Horizontal bars represent group means.

In the combination group, there were modest increases in the mean percentage of peripheral blood CD4^+^ regulatory T cells on and after day 15 in female monkeys that did not decline through day 72; similar changes in the CD4+ T reg profile were observed in male monkeys dosed with 50 mg/kg nivolumab, but these increases declined over time (Supplementary Fig. S14). Additionally, in the combination group, decreases in mean percentage naive CD4^+^ T cells and concomitant increases in central memory CD4^+^ T cells were noted during the recovery period. Statistically significant nonreversible increases in mean percentage splenic CD4^+^ regulatory T cells, nonreversible increases in mean percentage splenic CD25^+^FoxP3^+^CD8^+^ T cells, and decreases in mean percentage splenic naive CD8+ T cells, as well as concomitant increases in central memory CD8^+^ T cells during the recovery period (data not shown), were not considered different between groups dosed in combination or with 50 mg/kg nivolumab alone.

Collectively, these changes are consistent with the pharmacological mechanisms of action of relatlimab and nivolumab, and highlight the potential for enhanced effects of the antibodies when administered in combination.

## Discussion

Continuous exposure of T cells to cognate antigen leads to an exhausted, hyporesponsive phenotype characterized in part by the expression of multiple inhibitory receptors including PD-1, CTLA-4, LAG-3, and T-cell immunoglobulin and mucin domain-3 (30). LAG-3 has been shown to be expressed in TILs of several tumor types, including melanoma, hepatocellular carcinoma (HCC), and non-small cell lung cancer (NSCLC), often in parallel with increased PD-1 (31-33). Combination immunotherapies can result in improved clinical benefit compared with single checkpoint blockade, as exemplified by the combination of anti-PD-1, nivolumab, and anti-CTLA-4 (ipilimumab) approved in the US and other countries for multiple indications, including melanoma, NSCLC, renal cell carcinoma, HCC, colorectal cancer, and pleural mesothelioma (34,35). Targeting additional checkpoints, such as LAG-3, is a promising approach for overcoming resistance to ICB by targeting multiple inhibitory targets. LAG-3 acts in a nonredundant manner from that of PD-1 to suppress T-cell stimulation, and presents an exciting new opportunity for combination ICB that may improve clinical responses.

Preclinical data presented in recent years illustrate a clear synergy between the inhibitory receptors LAG-3 and PD-1 in controlling immune homeostasis, preventing autoimmunity, and enforcing tumor-induced tolerance (9,13). Combined treatment of mice with blocking antibodies against LAG-3 and PD-1 receptors resulted in more robust immune responses than either single-agent treatment (9,14-16). Similarly, in the *in vitro* primary T-cell superantigen stimulation assays reported here, only modest activity was observed from LAG-3 single-agent blockade with relatlimab compared with the substantially enhanced responsiveness in the context of co-blockade of LAG-3 and PD-1 with relatlimab and nivolumab, respectively. As expected for an IgG4 isotype antibody null for Fc receptor engagement, there was no measurable ADCC or complement-dependent cytotoxicity mediated by relatlimab. Toxicologic assessment of relatlimab in cynomolgus monkeys showed it to be generally well tolerated, alone and in combination with nivolumab. Relatlimab displayed no unexpected normal tissue cross-reactivity by IHC, except for observed on-target binding in the pituitary. The role of LAG-3 in the pituitary has not been determined.

Herein, we have demonstrated *in vitro* functional binding and blocking activity of relatlimab. The antibody binds with a higher affinity to activated human T cells than to activated cynomolgus T cells, most likely due to species differences in the sequence of the D1 domain that is targeted by the mAb. Relatlimab reversed the functional inhibition of T-cell activation by LAG-3 in an *in vitro* antigen-specific T-cell hybridoma assay, demonstrating that the antibody can functionally block the inhibition of T cells in the context of LAG-3 engagement by peptide-loaded MHC II.

Wang et al. recently reported the identification of FGL1 as a new putative ligand of LAG-3, presenting compelling evidence for its functional role in LAG-3 signaling and its potential clinical relevance in certain cancers (12). In our investigation of FGL1-mediated T-cell suppression, we showed that the engineered co-expression of a cell membrane-tethered FGL1 in the context of a mouse MHC II-positive APC resulted in more potent inhibition of T-cell responsiveness compared with engagement of the receptor by MHC II alone. In biochemical assays, relatlimab blocked the interaction of LAG-3 with both MHC II and FGL1. Consistent with these observations, relatlimab also strongly blocked the enhanced T-cell suppression observed in co-cultures with MHC II/FGL1 co-expressing APCs (**Fig. 4C**). In these assays, relatlimab restored T-cell cytokine production to a level equivalent to that observed for antibody treatment of T cells in co-culture with APCs expressing MHC II alone **(Fig. 4C)**. Additional work is needed to fully understand the nature of the interactions between these three molecules, but it is possible that a ternary complex of LAG-3, MHC II, and FGL1 could exist that may promote potent T-cell inhibition. Our data indicate that the FGL1/LAG-3 interaction is relatively weak (**Fig. 3C** and **D**) but, in the artificial context of APCs engineered for surface expression of FGL1, the increased valency likely fosters an avidity-driven enhancement of FGL1-mediated T-cell inhibition, as we observed. The mechanism by which soluble FGL1 can functionally engage LAG-3 in the periphery or intratumorally to inhibit T-cell responses remains unclear. FGL1 is produced by the liver and secreted into the bloodstream, where it plays a role in hepatocyte regeneration and metabolism to suppress environmentally induced inflammation (12,36-39). FGL1 has been reported to promote invasion and metastasis in gastric cancer and to mediate drug resistance in lung and liver cancer (40-42). It has been suggested that FGL1 may associate with extracellular matrix components to facilitate LAG-3 interactions, similar perhaps to the interactions of the latent transforming growth factor (TGF)-β complex with α_v_ integrins and the subsequent release of active TGF-β (43), but this has yet to be demonstrated. There is some evidence that FGL1 may form high molecular weight oligomers with FGL2 (44), but it is unclear whether FGL1 is able to assemble into homogeneous high molecular weight complexes alone. Dimeric Fc-tagged FGL1 protein, carefully processed to ensure a homogenous dimer preparation, failed to inhibit in the 3A9 T-cell hybridoma assay as previously reported (12), but we observed some evidence of 3A9 cell inhibition when cells were treated with hexameric FGL1. These differences in results may reflect differences in the biochemical properties of the purified proteins used in the different assays. It remains to be determined what the physical oligomerization state of FGL1 is in the periphery and the TME, and it remains unclear what effect soluble LAG-3 in either compartment may have on the ability of tumor-expressed FGL1 to functionally engage LAG-3 on T cells.

Biochemical and biophysical analyses demonstrated that the epitope of relatlimab resides in the insertion loop of the D1 domain of LAG-3, and is centered on the peptide, H63–W70. Wang et al. showed evidence that the Y77F mutation in the C’ strand of D1, which has been shown to abrogate MHC II interaction with LAG-3, did not perturb the interaction with FGL1 (12). These results support the hypothesis that FGL1 and MHC II interaction sites on LAG-3 are independent of one another. While the mechanism of soluble FGL1-mediated inhibition remains to be determined, our data demonstrate that relatlimab can block its interaction with LAG-3, supporting the potential utility of relatlimab in cancer indications where FGL1 expression is high (e.g. HCC), or where its expression correlates with poor prognosis (e.g. lung adenocarcinoma) (12,42).

The efficacy and manageable safety profile of relatlimab combined with nivolumab has been demonstrated in the phase II/III clinical trial RELATIVITY-047, where prolonged PFS benefit was observed with relatlimab combined with nivolumab compared with nivolumab monotherapy in patients with previously untreated metastatic or unresectable melanoma (17,18). Collectively, these data support the development of a combination of relatlimab plus nivolumab as a promising therapeutic strategy in clinical oncology that has the potential to enhance antitumor responses and broaden the range of responding cancer types compared with nivolumab monotherapy.

## Acknowledgments

Medical writing support and editorial assistance were provided by Katerina Pipili, PhD, and Matthew Weddig of Spark Medica Inc., funded by Bristol Myers Squibb, according to Good Publication Practice guidelines. Jun Zhang contributed to antibody generation efforts. Christine Bee and Mohan Srinivasan contributed to SPR studies with additional support and review from Gavin Dollinger and Andrew Drake. Winse Morishige assisted with SPR data visualization. Jennifer Price provided PCR study support and Olufemi Adelakun contributed to IHC studies. Rangan Vangipuram managed antibody purification work, and Sharon Viajar aided in protein purification. Peter S. Lee provided guidance on structural characterization, and Arvind Rajpal provided input on biophysical studies. In support of the 1-month monkey toxicity study, Jagannatha Mysore provided pathology analysis, Carol Gleason provided biostatistical analysis, and Gautham Rao provided input on study design and analyses of pharmacodynamic endpoints. The authors particularly thank Dario Vignali for discussions, feedback, and support as a subject matter expert of LAG-3 biology, as well as for providing the 3A9 hybridoma cell lines and C9B7W antibody used in these studies.

