## Supplement for "Preclinical Characterization of Relatlimab, a Human LAG-3–Blocking Antibody, Alone or in Combination With Nivolumab"

### Supplementary Methods

#### Relatlimab epitope characterization

##### Binding affinity of relatlimab to human LAG-3 by surface plasma resonance

The avidity binding measurement was determined by capturing relatlimab on anti-κ monoclonal antibody (Southern Biotech, Birmingham, AL, USA)-coated CM5 sensor chip surfaces (4.5–8.5 kRU; Biacore, Uppsala, Sweden). HBS-EP (GE Healthcare, Chicago, IL, USA) was used as a running buffer and for making dilutions. Multiple concentrations (4–16 nM) of human lymphocyte-activation gene 3 (LAG-3) fused with the Fc of human immunoglobulin (Ig) G1 (hLAG3–hFc) (R&D Systems, Minneapolis, MN, USA) were flowed over the captured monoclonal antibody for 5 minutes followed by 20 minutes of dissociation at 25ºC. Each cycle was regenerated with 50 mM HCl/75 mM NaCl and 50 mM NaOH followed by HBS-EP wash. The kinetic data were generated using BIAevaluation version 3.2.

The monovalent binding affinity was determined by capturing hLAG3-hFc (R&D Systems) for 30, 60, or 90 seconds in separate flow cells over a high-density anti-human IgG Fc polyclonal antibody (Southern Biotech)-coated chip surface to achieve ligand loading levels of approximately 30, 80, and 100 RUs, respectively. Binding measurements and analyte dilutions were made at either pH 6.0 or pH 7.4 using the running/dilution buffers (10 mM Bis-Tris pH 6.0, 150 mM NaCl, 0.05% [v/v] TWEEN^®^20, 1 g/L bovine serum albumin [BSA]) and (10 mM HEPES pH 7.4, 150 mM NaCl, 0.05% [v/v] TWEEN^®^20, 1 g/L BSA), respectively. Multiple concentrations (250 pM–20 nM, three-fold dilution series) of relatlimab antigen-binding fragment (Fab) 3.5 analyte were flowed over the captured human LAG3-Fc for 2 minutes followed by 5 minutes of dissociation at 37ºC. Each cycle was regenerated with a 20-second pulse of 73 mM phosphoric acid at 30 μL/minute. The kinetic data were generated using Biacore T200 BIAEvaluation software.

##### Antigen binding and epitope specificity

The binding of relatlimab to recombinant hLAG-3–mFc fusion protein and synthetic human LAG-3 (hLAG-3) insertion-loop peptides was determined using enzyme-linked immunosorbent assay (ELISA). The custom peptides consisted of the full-length loop human LAG-3 peptide, corresponding to residues G48-Y77 of the mature LAG-3 polypeptide (GPPAAAPGHPLAPGPHPAAPSSWGPRPRRY; NP_002277.4), or of a set of 10 synthetic overlapping peptides (12-mer peptides overlapping by 10 residues spanning the hLAG-3 peptide sequence; BioMol, Intl., Exeter, UK). Serial dilutions of relatlimab were tested by ELISA for binding to microtiter plate-immobilized LAG-3–mFc fusion protein (BMS, NJ, USA) or peptide, and detected using peroxidase-conjugated secondary reagent (Jackson ImmunoResearch, West Grove, PA, USA).

**Carbene chemical footprinting of hLAG-3 and complexed with relatlimab Fab**

Human LAG-3 D1–D2 domains containing a His_6_ tag and human LAG-3 D1–D2 domains-His_6_ tagged protein complexed with relatlimab Fab (1:1.1 ratio) were subjected to differential carbene chemical labeling using the diazirine-containing reagent 4-(3-(Trifluoromethyl)-3H-diazirin-3-yl)benzoic acid. All samples were in 10 mM sodium phosphate and 50 mM NaCl, pH 7.4. Triplicate samples of the hLAG-3 and hLAG-3:relatlimab complex were mixed 1:1 with 20 mM 4-(3-(Trifluoromethyl)-3H-diazirin-3-yl)benzoic acid, incubated at room temperature for 30 minutes, and snap-frozen in liquid nitrogen. The frozen samples were irradiated for 20 minutes in a Suntest CPS+ light box with a xenon arc lamp filtered to pass 315–800 nm light (Atlas Material Testing Technology, Mount Prospect, IL, USA). The labeled samples were reduced, alkylated with iodoacetamide, and digested with chymotrypsin at 37C for 2.5 hours. The generated peptides were analyzed by liquid chromatography with tandem mass spectrometry (LC-MS/MS) on a Fusion Lumos mass spectrometer (ThermoFisher, San Jose, CA, USA). The LC-MS/MS data was searched using Byonic (Protein Metrics Inc., Cupertino, CA, USA) against the hLAG-3 D1–D2 construct with the following modifications: methionine and tryptophan oxidation, cysteine carbamidomethylation, carbene modification (+202.024164) to any amino acid, N-terminal glutamine to pyro-glutamic acid, asparagine deamidation, and N-linked glycosylation of asparagine. Differential peptide level labeling was determined by comparing the integrated peak area ratios of each hLAG-3 peptide with the singly carbene-labeled peptide for both the hLAG-3 and hLAG-3:relatlimab complex samples. Statistically significant peptides (*P* < 0.01) were determined using Student’s t-test. Amino acid residue level labeling was determined using the method described by Jumper et al. (1). All the tandem mass spectrometry (MS/MS) scans associated with a singly carbene-labeled peptide were summed into a single MS/MS spectrum, and then the ratio of peptide MS/MS fragment ion intensity for each fragment ion containing or lacking the presence of the carbene label was determined for either a b- or y-ion series. Individual residue labeling ratios were identified by subtracting the residue level ratio from the (n-1) fragment from the residue level ratio (n) fragment and adjusting the resulting value by the peptide level ratio determined above. Any missing fragment ions were skipped and combined with the next identified fragment ion in the series. The summed residue level labeling should equal the peptide level labeling. Statistically significant residues (*P* < 0.01) were determined using Student’s t-test.

#### Fab 3.1 protein purification for X-ray crystallographic analysis

DNA constructs coding for human Fab of antibody LAG-3.1 (Fab 3.1) were cloned into pTT5 expression plasmids (GenScript, Piscataway, NJ, USA) with an osteonectin signal peptide on the N-terminus and a poly-histidine tag on the C-terminus of the heavy chain. Fab 3.1 was expressed in Expi293 cells, purified from clarified supernatant using Ni Sepharose^®^ Excel (GE Healthcare, Chicago, IL, USA), polished on a Superdex^®^ 200 16/60 column in 100 mM NaCl, 20 mM Tris pH 8.0, and concentrated to 20 mg/mL. For complex formation, excess peptide (final concentration of 1 mM) was added to Fab 3.1.

#### Crystallographic analysis of hLAG-3 epitope in complex with relatlimab by x-ray diffraction

A 16-mer peptide corresponding to residues within the D1 loop (Ac-PGPHPAAPSSWGPRPR-amide) of hLAG-3 was synthesized at Elim Biopharmaceuticals, Inc. (Hayward, CA, USA) to >98% purity. The Fab 3.1 solution was mixed 1:1 with 0.1 M Bis-Tris, pH 5.5; 25% PEG 3350 in a total volume of 0.4 µL, and peptide crystals were grown by sitting drop vapor diffusion at 20°C using the same buffer as the mother liquor. X-ray diffraction data were collected at beamline 17ID (wavelength 1.0 Å) at the Advanced Photon Source (Argonne, IL, USA). Complex diffraction data were collected at 2.4 Å resolution. The Fab 3.1–peptide complex was determined by molecular replacement with Phaser using models for the constant and variable domains. The Fab model was iteratively built using Coot and refined in PHENIX (2,3). The peptide was built *de novo* into the model using Coot and refined in PHENIX. In the final structure, 96.2% of the residues are in favored regions of the Ramachandran plot with 0.5% outliers, as calculated by MolProbity. X-ray diffraction data collection and refinement statistics are reported in **Supplementary** **Table S6**. Kabat numbering was applied to the variable domains of the Fabs.

#### Relatlimab binding to hLAG-3 and cynomolgus monkey LAG-3 by flow cytometry

Serial dilutions of relatlimab were tested for binding to engineered Chinese hamster ovarian cell line-S (CHO-S) cells expressing hLAG-3 or cynomolgus monkey LAG-3, and to activated human CD4+ or cynomolgus T cells using flow cytometry. Bound relatlimab was detected using a phycoerythrin (PE)-conjugated secondary antibody specific to human Fcγ (PE-Gt-F(ab’)2-anti-hFcγ, Jackson ImmunoResearch). Cells were analyzed on FACSCanto or LSR II flow cytometers running BD FACSDiva™ acquisition control software (BD Biosciences, Franklin Lakes, NJ, USA). Post-acquisition analysis was performed using FlowJo software v4.4.3 (Tree Star, Inc., Ashland, OR, USA).

#### Blockade of LAG-3–Fc binding to MHC II+ Daudi B cells

The potency of relatlimab in blocking the interaction of LAG-3 and MHC II cells was evaluated by means of a flow cytometry blocking assay. Serially diluted relatlimab or isotype control antibody was mixed with a fixed concentration (approximately effective concentration [EC]_80_ for binding to Daudi cells) of human LAG-3 extracellular domain (domains D1-D4) fused with the Fc of mouse IgG1 (hLAG-3–mFc) antigen in solution and allowed to precomplex for 20 minutes prior to adding to human MHC II+ Daudi B lymphoma cells (ATCC, Manassas, VA, USA). Conjugated secondary antibody specific to mouse Fcγ (PE-Gt-F(ab’)2-anti-muFcγ, Jackson ImmunoResearch) was used to detect the presence of bound hLAG-3–mFc antigen. The presence of cell-bound LAG-3-mFc antigen was analyzed on a FACSCanto flow cytometer (BD Biosciences, San Jose, CA, USA) in conjunction with software packages FlowJo (V. 10, FlowJo, Ashland, OR, USA) and Prism (V. 5.01, Graphpad Software, La Jolla, CA, USA).

#### Octet^®^ biolayer interferometry assays of LAG-3/MHC II interaction and blockade by relatlimab

Recombinant human leukocyte antigen-DR isotype (HLA-DR) with a C-terminal biotin modification was generated at Bristol Myers Squibb using human *HLA-DRA* and *HLA-DRB1* genes cloned into pTT5 expression plasmids (GenScript, Piscataway, NJ, USA), which were subsequently co-expressed in Expi293 cells, purified from clarified supernatant using Ni Sepharose^®^ Excel (GE Healthcare, Chicago, IL, USA), and polished on a Superdex^®^ 200 16/60 column in phosphate buffered saline (PBS). The purified heterodimer was subsequently biotinylated using a BirA biotin-protein ligase bulk reaction kit (Avidity, Aurora, CO, USA), and buffer exchanged into PBS for further experiments. A recombinant hLAG-3 (D1–D4)–hFc fusion protein was produced by cloning hLAG-3 (D1–D4) into a pTT5 expression plasmid (GenScript, Piscataway, NJ, USA) with an osteonectin signal peptide on the N-terminus and a C-terminal hFcG1 tag. Clarified supernatants were purified using MabSelect SuRe™ LX resin (GE Healthcare), washed with PBS, eluted in 100 mM pH 3.6 sodium citrate buffer, and neutralized with 1 M pH 8.0 Tris buffer. hLAG-3 (D1–D4) was further purified on a Superdex^®^ 200 column into 1× PBS. HLA-DR1 was captured at 10 µg/mL for 180 seconds in PBS 0.05% BSA with 0.05% TWEEN^®^20 pH 7.4 (Sigma-Aldrich, St Louis, MO, USA; PBST-BSA), followed by a 60-second wash with PBST-BSA. hLAG-3 (D1–D4)–hFc (1 µM) was tested for binding to the captured HLA-DR protein over 300 seconds.

##### Octet^®^ biolayer interferometry assays of LAG-3/FGL1 interaction and blockade by relatlimab

The binding of recombinant human fibrinogen-like protein-1 (FGL1)-mFc fusion protein and recombinant human FGL1-FD-mFc fusion protein (consisting of only the fibrinogen domain of human FGL1 [residues 74–312] with an N-terminal mouse Fc tag and produced by BMS) to recombinant hLAG-3–hFc was investigated on an Octet^®^ HTX instrument with anti-human IgG Fc capture (AHC) biosensors (ForteBio). LAG-3–hFc was captured at 10 µg/mL for 300 seconds in PBST-BSA, followed by a 60-second wash with PBST-BSA. Serial dilutions of FGL1-mFc and FGL1-FD-mFc were tested for binding to the captured LAG-3–hFc fusion protein. To assess the blockade of LAG-3/FGL1 engagement by relatlimab, LAG-3–hFc was first captured on AHC biosensors at 10 μg/mL for 600 seconds in PBST-BSA, followed by a 30-second wash with PBST-BSA. The biosensor was next quenched with a cocktail of keyhole limpet hemocyanin (KLH) human IgG1 (hIgG1), KLH-hIgG2, and KLH-hIgG4 antibodies for 600 seconds in PBST-BSA to block the surface. This was followed by a 30-second wash with PBST-BSA, exposure to buffer alone or relatlimab (200 nM) in PBST-BSA for 1000 seconds, and a 10-second wash with PBST-BSA. Finally, for biosensors exposed to buffer alone, FGL1-FD-mFc (2 μM) in the absence of relatlimab was tested for binding. For biosensors previously exposed to relatlimab, FGL1-FD-mFc (2 μM) in the presence of excess relatlimab (200 nM) was tested for binding. For all biolayer interferometry assays, the temperature was 30°C with a shake speed of 1000 RPM. Data were visualized on ForteBio data analysis software.

#### Functional cell-based bioassay of LAG-3/MHC II interaction and blockade by relatlimab

Functional analysis of LAG-3 antibody functional potency was performed using a mouse T-cell hybridoma assay (4). Mouse 3A9 T cells expressing full-length hLAG-3 and a T-cell receptor responsive to peptide (48-DGSTDYGILQINSRW-62) of hen egg lysozyme and LK35 cells presenting matched MHC II (I-A^k^) were dispensed into wells of a microtiter plate at a 2:1 ratio, in the presence of 1) titrated peptide (1:3 from 30 uM) and antibody at a fixed, saturating concentration (20 ug/mL), or 2) titrated antibody (1:3 from 20 ug/mL) and peptide at a fixed, suboptimal concentration (150 nM). Supernatants were collected after 15 hours of culture for measurement of secreted mouse interleukin (IL)-2 cytokine by the 3A9 T cells by ELISA (BD Biosciences, cat. number 55148). Following the addition of substrate, assay plates were read on a plate spectrophotometer at 650–570 nm.

#### Functional cell-based bioassay of LAG-3/FGL1 interaction and blockade by relatlimab

To investigate the inhibitory effect of LAG-3 engagement by FGL1, a modified version of the 3A9 functional screening assay was developed. An expression vector was prepared that appended human FGL1 (including the coiled-coil and fibrinogen domains) to the N-terminus of a cDNA sequence corresponding to the intracellular and transmembrane domains of the human FIBCD1 type II transmembrane protein. This construct was transfected into the LK35.2 antigen-presenting cell line to stimulate 3A9-hLAG-3 cells as described above. The resulting modified antigen-presenting cell line expresses both mouse MHC II and a membrane-tethered form of human FGL1.

#### Antibody-dependent cellular cytotoxicity

Peripheral blood mononuclear cells (PBMC) effector cells were purified from heparinized whole blood samples from two donors by standard Ficoll-Paque (Mediatech, Manassas, VA, USA) separation and cultured overnight in the presence of IL-2 (R&D Systems). Activated T cells from three human donors, activated for 72 hours with anti-CD3 and anti-CD28 antibodies (clones UCHT1 and CD28.2, respectively; BD Biosciences, NJ, USA), were labeled with 20 mM BATDA (PerkinElmer, Waltham, MA, USA) and dispensed at 10,000 cells/well into the assay plate to result in a final target to effector cell ratio of 1:50. Five-fold serial dilutions of relatlimab, positive control non-fucosylated antibody (human anti-CD30-NF), and isotype control antibodies (hIgG4 and hIgG1-NF) were added to effector/target cell mixtures to result in final antibody concentrations of 10 μg/mL to 3.2 ng/mL. After 1 hour of incubation at 37°C, assay supernatants were diluted in europium solution (PerkinElmer) in a flat-bottom 96-well plate. The resulting reaction was read with a Fusion-Alpha TRF Reader using a 400-μsecond delay, and 330/80 excitation and 620/10 emission filters. Target cells co-cultured with effector cells in absence of antibody provided the control for background antibody-independent lysis, while target cells co-cultured with effector cells then lysed with methanol represented maximal release in the assay. Antibody-dependent, percent-specific lysis was calculated based on counts per second (CPS) with the following formula: [(test CPS – mean background)/(mean maximum CPS – mean background)] × 100 where background is effector cells + target cells with no antibody, and maximal lysis is effector cells + target cells + methanol. Percent-specific antibody-dependent lysis was plotted against antibody concentration, and nonlinear regression, sigmoidal dose–response analysis was used to calculate the effective concentration needed to achieve a 50% maximal response (EC_50_) values for each assay in Graphpad Prism software.

#### Antitumor activity of anti–LAG-3 and anti–PD-1 in mouse models

##### Mice

Female C57BL/6 mice were obtained from Charles River Laboratories (Wilmington, MA, USA) and female A/J mice were obtained from Harlan (Dublin, VA, USA). Throughout the study all animals were provided chow (Prolab^®^ Isopro^®^; Dean’s Animal Feeds, Redwood City, CA, USA) and water ad libitum. The study protocol was approved by the Institutional Animal Care and Use Committee prior to study initiation, and all animal husbandry was performed according to Medarex Standard Operating Procedures. The Medarex Animal Facility is accredited by the Association for Assessment and Accreditation of Laboratory Animal Care.

##### Cell culture and syngeneic tumor studies

MC38 colon adenocarcinoma and SA1/N fibrosarcoma tumor cells from the BMS Master Cell Bank (Princeton, NJ, USA) were thawed and maintained in Dulbecco’s modified Eagle’s medium (Cellgro) supplemented with 10% fetal bovine serum (FBS; Gemini Bio-Products, West Sacramento, CA, USA). Cells were validated to be free of adventitious agents by polymerase chain reaction (PCR; Idexx Bioanalytics, Columbia, MO, USA). Cells were harvested near 80% confluence, washed in FBS-free medium three times, and resuspended in PBS to provide subcutaneous injections of 1.0E+06 cells (0.1 mL) into the right flank of each study animal. Tumor volumes were measured with an electronic caliper (l × w × h/2) and animals were randomized into study groups with similar tumor size ranges. Mice with palpable tumors on day 7 post-tumor implantation (average volume 75–100 mm^3^) were injected interperitoneally at a dose of 10 mg/kg (0.2 mg) for all antibodies, as appropriate by group, on days 7, 10, and 14. Tumor size and body weight of study animals were monitored and recorded twice weekly until tumors had reached endpoint (≤1500 mm^3^).

##### Antibodies

###### Anti-mouse PD-1 monoclonal antibody (4H2)

Chimeric surrogate antibody 4H2 was as described previously (5).

###### Anti-mouse LAG-3 monoclonal antibody (C9B7W)

Surrogate antibody C9B7W was as described previously (4).

###### Anti-mouse LAG-3 monoclonal antibody (19C7)

To obtain 19C7, rats were immunized with mouse LAG-3–Ig, followed by splenocyte fusion and screening of clones for reactivity to the receptor by ELISA. The 19C7 antibody was selected by virtue of its high affinity for mouse LAG-3 and for its ability to block the interaction between murine LAG-3–Ig and MHC Class II expressed on murine A20 lymphoma cells. After the variable (V) region sequences of 19C7 were determined, the variable light chain (Vl) and variable heavy chain (Vh) sequences were grafted onto the murine IgG1 Fc region containing a D265A mutation for reduced Fc receptor engagement. The chimeric antibody was then expressed from a transfected CHO cell line.

###### Mouse IgG1 isotype control antibody

Purified mouse IgG1 (MOPC-21) was obtained from Bio X Cell (West Lebanon, NH, USA).

#### Immunohistochemistry analysis in normal human tissues

Immunohistochemistry analyses were performed in a selected panel of normal human tissues. Fresh frozen normal human tissues were purchased from commercial tissue networks/vendors (Analytical Biological Services, Inc., Wilmington, DE, USA; Asterand, Inc., Detroit, MI, USA; and Cooperative Human Tissue Network, Philadelphia, PA, USA). Stained slides were evaluated under a light microscope. To detect tissue binding, the fluoresceinated forms of relatlimab, LAG3.1-G4P, and isotype control antibody were prepared in-house and designated as relatlimab-FITC, LAG3.1-G4P-FITC, and huIgG4-FITC, respectively. Cryostat sections at 5 µm were fixed with acetone and then washed twice with PBS at room temperature. LAG-3–expressing lymphocytes and LAG-3 negative elements in tonsil sections were used as positive and negative control tissues respectively. In addition, cryostat sections of Chinese hamster ovary (CHO)-S-human LAG-3 cells and CHO-S cells were also used as positive and negative controls. Endogenous peroxidase activity was blocked by incubation with peroxidase block supplied in Dako EnVision System (Agilent, Carpinteria, CA, USA; Catalog K4011). Slides were washed in PBS then incubated with Dako protein block supplemented with 0.5% human γ globulins to block the nonspecific binding sites. Subsequently, primary antibodies (relatlimab-FITC or LAG-3.1-G4P-FITC) or isotype control (hIgG4-FITC) were applied onto sections and incubated for 1 hour. After washing, slides were subjected to serial 30-minute incubations separated by repeat washes, with rabbit anti-FITC antibody, peroxidase-conjugated anti-rabbit IgG polymer, and finally diaminobenzidine (DAB) substrate-chromogen solution (Dako EnVision System; Agilent, Santa Clara, CA, USA). Slides were washed with deionized water, counterstained with Mayer’s hematoxylin, dehydrated, cleared, and coverslipped with Permount following routine histological procedures.

To verify the cell type within the pituitary gland to which relatlimab bound, double staining of LAG-3.1-G4P-FITC with five pituitary hormone antibodies on cryostat sections was performed. In the same manner as above, slides were incubated with Dako protein block supplemented with 1% human γ globulins to block nonspecific binding sites. To reduce background further, the PBS used in immunofluorescence contained high salt and low detergent (300 mM NaCl and 0.01% TWEEN^®^20 ). Subsequently, LAG-3.1-G4P-FITC with each of the following commercial mouse monoclonal antibodies anti-adrenocorticotropic hormone (Dako) and anti-thyroid stimulating hormone (Abcam, Cambridge, MA, USA), or rabbit pAbs anti-growth hormone (Dako), anti-prolactin (Dako), and anti-luteinizing hormone (Dako) were simultaneously applied onto sections. Human IgG4-FITC, with either mouse IgG1 or rabbit IgG, was used as isotype control. After washing, slides were incubated with secondary antibodies: alexa Fluor 488–conjugated goat anti-FITC/Oregon-Green (Invitrogen), Cy3-labeled donkey anti-mouse (Jackson ImmunoResearch), and Cy3-conjugated donkey anti-rabbit IgG (Jackson ImmunoResearch). After washing, sections were counterstained with Hoechst 33342 (Invitrogen) mounting with ProLong-Gold Antifade reagent (Invitrogen). To assess the specificity of the LAG-3.1-G4P-FITC staining, pre-absorption of the primary antibody with human LAG-3 fusion protein was performed by pre-incubating LAG-3.1-G4P-FITC with 5- or 10-fold excess of human LAG-3 for 2 hours at room temperature before applying to the sections from two pituitary samples following respective protocols for immunoperoxidase and immunofluorescent methods. PBS supplemented with 0.5% human γ globulins and 0.5% BSA was used as a diluent for both primary and secondary antibodies. For more stringent nonspecific blocking conditions, Dako protein block supplemented with 1% human γ globulins or PBS supplemented with 1% human γ globulins was used as a blocking buffer or diluent, respectively. Both the staining intensity and frequency were evaluated by light microscopy.

#### PCR for pituitary expression of LAG-3

PCR was performed to confirm LAG-3 mRNA expression in both CD8+ T cells (positive control) and in pituitary tissue RNA samples. Nuclease-free water was used as the no template control. LAG-3 primers and housekeeping gene *PPIA* (Cyclophilin A) primers were designed using Primer Blast software (http://www.ncbi.nlm.nih.gov/tools/primer-blast/). LAG-3 primer sets and expected PCR amplicon sizes were based on GenBank Accession No. NM_002286.5. Primer sets F38-R333, F732-R1365, and F732-R1664 were used in conventional PCR; primer set F732-R840 was used in real-time PCR. RNA was quantitated and then reverse transcribed to cDNA (SuperScript II; Invitrogen, Carlsbad, CA, USA). For PCR, cDNA (50 ng) was mixed with the specific primer pairs and amplified using the Platinum R TaqDNA Polymerase High Fidelity kit (Invitrogen) using the thermal cycling profile: 95°C, 5 minutes; 95°C, 30 seconds; 65°C, 30 seconds; 72°C, 4 minutes (40 cycles); and 72°C, 7 minutes.

#### Toxicity studies in cynomolgus monkey model

For 1-month and 3-month toxicity studies, cynomolgus monkeys (M. fascicularis) of Mauritian origin were obtained from Charles River BRF, Inc. (Houston, TX, USA) and Buckshire Corporation (Perkasie, PA, USA). Male and female monkeys (3.5 to 4.6 and 3.4 to 5.4 years old, respectively, and weighing between 3.9 and 5.4 kg, and 3.0 and 4.1 kg, respectively) were randomly assigned by algorithm to study groups (Study Log, South San Francisco, CA, USA). Monkeys were housed under appropriate environmental (temperature, humidity, light cycle) conditions and were offered purified and chlorinated tap water ad libitum and fed Diet #2050-C Protein Primate Diet (Harlan) with a daily ration of biscuits. The study procedures were approved by the Institutional Animal Care and Use Committee, and the monkeys were monitored daily for clinical signs including body weight, cardiovascular and ophthalmologic evaluations, and veterinary physical examinations. For the studies, animals were dosed intravenously with the indicated antibody test articles at the doses and schedules summarized in **Supplementary Table S1** and **S2**. Animals were administered a single intramuscular bolus of KLH (Calbiochem/Sigma-Aldrich, St. Louis, MO) (dosed day 1, 1-month study; day 57, 3-month study) to evaluate drug-related immunological responses. Blood samples for evaluation of KLH-specific antibody responses were collected at various timepoints. In the 1-month study, scheduled necropsies were conducted after the fifth weekly dose (3/sex/group) and following a 6-week recovery period (2/sex/group, if available). In the 3-month study, scheduled necropsies were conducted on 24 monkeys (4/sex/group) after the 13th weekly dose while the remaining 12 monkeys (2/sex/group) were necropsied after a 10-week recovery period.

##### Cynomolgus monkey splenic T-lymphocyte subset phenotyping

At scheduled necropsies, unfixed sections were collected from the same region of the spleen using aseptic methods, weighed, minced, and placed on wet ice into tubes containing 1× Hanks balanced salt solution buffer for analysis of splenic T-lymphocyte subsets by flow cytometric methods. Samples were analyzed by flow cytometry for the following populations: total T cells (CD45+CD16-CD3+), helper T cells (CD45+CD16-CD3+CD4+CD8-), and cytotoxic T cells (CD45+CD16-CD3+CD8+CD4-). Samples were reported as percentage of splenic lymphocytes. In addition, samples were also analyzed for the following populations: CD4 regulatory T cells (CD3+CD4+CD8-CD25+Foxp3+), naïve CD4 T cells (CD3+CD4+CD8-Foxp3-CD95-CD28+), central memory CD4 T cells (CD3+CD4+CD8-CD95+CD28+), effector memory CD4 T cells (CD3+CD4+CD8-CD95+CD28-), naïve CD8 T cells (CD3+CD8+CD4-CD95-CD28+), central memory CD8 T cells (CD3+CD8+CD4-CD95+CD28+), effector memory CD8 T cells (CD3+CD8+CD4-CD95+CD28-), CD25+ activated CD4 T cells (CD3+CD4+CD8-CD25+), HLA-DR+ activated CD4 T cells (CD3+CD4+CD8-HLA-DR+), CD25+ activated CD8 T cells (CD3+CD8+CD4-CD25+), HLA-DR+ activated CD8 T cells (CD3+CD8+CD4-HLA-DR+), and CD25+FoxP3+ CD8 T cells (CD3+CD8+CD4-CD25+Foxp3+), and reported as percentages of respective T-cell subset populations (either CD4+CD8- or CD8+CD4-).

##### Ex vivo recall responses to KLH

Whole blood samples were collected from all available monkeys into sodium heparin collection tubes once prior to study start, on day 22 pre-dose, and on day 57 pre-dose for assessment of *ex vivo* recall responses to KLH. Samples were mixed gently and kept at ambient temperature prior to analysis for intracellular cytokines and T-cell activation markers. Briefly, PBMCs isolated from whole blood were activated with KLH or anti-monkey CD3+ control with anti-CD28 and anti-CD49d costimulatory antibodies) followed by treatment with brefeldin A for approximately 16 to 20 hours. Following *ex vivo* culture, PBMCs were surface stained for expression of CD3, CD4, and CD8, along with a viability marker. After fixing and permeabilizing, the cells were labeled with anti–interferon-γ (IFN-γ), anti–tumor necrosis factor-α (TNF-α), and anti-CD69 fluorophore-conjugated antibodies, and analyzed by flow cytometry for the following CD4+CD8- and CD8+CD4- T-cell subsets: IFN-γ+, TNF-α+, CD69+, CD69+IFN-γ+, CD69+TNF-α+, and CD69+TNF-α+IFN-γ+. Results were reported as percentages of respective T-cell subset populations (either CD4+CD8- or CD8+CD4-). Only data following KLH stimulation were reported.

### Supplementary Tables

Table S1. Experimental design of 4-week multi-dose toxicity^a^ study in cynomolgus monkeys

| **Group number** | **Dosing/ route** | **Relatlimab (mL/kg)** | | | **Nivolumab (mL/kg)** | | | **Number and gender of monkeys** |
| --- | --- | --- | --- | --- | --- | --- | --- | --- |
| 1 | QW^b^/  IV | **0** | 0 | 5^d^ | **0** | 0 | 5^e^ | 5 M, 5 F |
| 2 |  | **30** | 20^c^ | 1.5 | **0** | 0 | 5^e^ | 5 M, 5 F |
| 3 |  | **100** | 20^c^ | 5 | **0** | 0 | 5^e^ | 5 M, 5 F |
| 4 |  | **0** | 0 | 5^d^ | **50** | 10^c^ | 5 | 5 M, 5 F |
| 5 |  | **100** | 20^c^ | 5 | **50** | 10^c^ | 5 | 5 M, 5 F |

^a^Clinical evaluation parameters included survival, clinical observations, body weight, and qualitative feeding behavior.

^b^Doses administered as slow intravenous (IV) infusions (1.2 to 2 mL/min) once weekly (QW) for 1 month (five total doses administered).

^c^For dosing purposes, concentrations of relatlimab and nivolumab were rounded.

^d^Vehicle for relatlimab: 10 mM sodium citrate, 10 mm phosphate buffer, 150 mm NaCl, 0.05% Tween 80, pH 5.5 ± 0.2.

^e^Vehicle for nivolumab: 20 mM sodium citrate, 50 mM NaCl, 3.0% mannitol, 20 µM diethylenetriamine pentaacetate, 0.02% polysorbate 80. pH 6.0.

Table S2. Experimental design of 3-month multi-dose toxicity study^a,b^ in cynomolgus monkeys

| **Group number/ therapy** | **Dose level (mg/kg/dose)** | **Dosing route^c^** | **Dose volume (mL/kg)** | **Dose rate (mL/kg/hr)** | **Conc.**  **(mg/mL)** | **Number of animals** | |
| --- | --- | --- | --- | --- | --- | --- | --- |
|  |  |  |  |  |  | **Dosing period^d^** | **Recovery period^e^** |
| 1/Vehicle control | 0 | IV | 10 | 30 | 0 | 4 M, 4 F | 2 M, 2 F |
| 2/Relatlimab | 30 | IV | 10 | 30 | 3 | 4 M, 4 F | 2 M, 2 F |
| 3/Relatlimab | 100 | IV | 10 | 30 | 10 | 4 M, 4 F | 2 M, 2 F |

^a^Clinical evaluation parameters included survival, toxicokinetics, clinical observations (including physical examinations), body weight, food evaluation, ophthalmologic examination, electrocardiology, clinical pathology (hematology, coagulation, clinical chemistry, and urinalysis), immunogenicity, T-cell–dependent antibody responses, immunophenotyping (peripheral blood and splenic immunophenotyping), *ex vivo* recall response assessment, organ weight, and gross and microscopic pathology analyses.

^b^The no-observed-adverse-effect level was considered to be 100 mg/kg/week (combined sex mean area under the curve[0-168h] 1,180,000 µg•h/mL).

^c^Dose formulations were administered to the appropriate animals by 20-minute intravenous infusion via brachial or saphenous vein once weekly for 3 months (13 doses).

^d^Animals were necropsied on day 92 at the end of the 3-month dosing period, with the last dose administered on day 85.

^e^Animals were necropsied at the end of the 10-week recovery period on day 148.

Table S3. X-ray diffraction data collection and refinement statistics

| **Data collection** |  |
| --- | --- |
| Beamline | APS 17-ID |
| Resolution (Å) | 83.855–2.398 |
| Space group | P 21 21 21 |
| Unit cell dimensions |  |
| a, b, c (Å) | 64.238 112.62 125.619 |
| α, β, γ (°) | 90, 90, 90 |
| *R*_merge_ | 0.194 (0.883) |
| *I*/σ(*I*) | 6.0 (1.7) |
| *Pearson’s CC*_1/2_ | 0.84 (0.76) |
| Completeness (%) | 99.51 (96.0) |
| Redundancy | 6.3 (5.7) |
| **Refinement** |  |
| Resolution | 56.310–2.398 |
| Number of reflections | 36287 |
| *R*_work_ / *R*_free_ (%) | 22.1 / 27.0 |
| Number of atoms |  |
| Protein | 6688 |
| Ligand | 0 |
| Water | 84 |
| Average B factors |  |
| Protein | 50 |
| Root mean square deviations |  |
| Bond lengths (Å) | 0.002 |
| Bond angles (°) | 0.516 |
| Ramachandran (%) |  |
| Favored | 96.16 |
| Allowed | 3.37 |
| Outliers | 0.47 |
| **PDB code** | XXXX^a^ |

^a^Structures to be deposited in the RCSB Protein Data Bank.

CC, correlation coefficient; R, reliability value.

Table S4. Tissue binding properties of relatlimab

| **Tissue types** | **Relatlimab-FITC** |
| --- | --- |
| CHO-S/hLAG-3 cells | 3+, v-fre |
| Tonsil #1   - Lymphocytes - Other | 3+, rare  -, ± |
| Tonsil #2   - Lymphocytes - Other | 3+ rare  - |
| Spleen   - White pulp - Red pulp - Other | 2+, e-rare, lymphocytes  1+, oca, MNC  - |
| Thymus | - |
| Pituitary #1   - Adenohypophysis - Neurohypophysis - Other | 3+, rare-oca  -  - |
| Pituitary #2   - Adenohypophysis - Neurohypophysis - Other | 2+, 3+, oca  -  -, ± |
| Pancreas   - Acini - Islets - Other | -  -  ±. 1+, oca, interstitial matrix |
| Cerebrum | - |
| Cerebellum | - |
| Heart | - |
| Liver | - |
| Lung | - |
| Kidney | - |

Grading scale: -, negative; ±, equivocal; 1+, weak; 2+, moderate; 3+, strong; 4+, intense. E-rare, extremely rare <1%; rare, 1 ~ <5%; rare~oca, rare~occasional, 5 ~ <10%; oca, occasional, 10 ~ <25%; oca~fre, occasional~frequent, 25 ~ <50%; fre, frequent, 50 ~ <75%; v-fre, very frequent, >75%; MNC, mononuclear cell; PALS, periarterial lymphatic sheath.

Table S5. Histopathology findings in cynomolgus monkeys treated with relatlimab and nivolumab in combination (Group 5) or nivolumab alone (Group 4) from a 4-week repeat-dose toxicity study

| **End of dose** | **End of recovery** | **Additional comments** |
| --- | --- | --- |
| **Group 5** | |  |
| **Scheduled necropsy on day 30** | **Scheduled necropsy on day 72** | - mean relatlimab AUC[0-168h] 514,000 µg•h/mL - Histopathological findings in monkey 5103 included slight lymphoplasmacytic inflammation of the choroid plexus; minimalto-moderate lymphohistiocytic inflammation of the vasculature of the brain parenchyma, meninges, and spinal cord; and minimal-to-moderate mixed-cell inflammation of the epididymis, seminal vesicles, and testes |
| Lymphoplasmacytic inflammation of the choroid plexus   - 3 of 3 males (Grade 1) and 2 of 3 females (Grade 1 or 2) | Lymphoplasmacytic inflammation of the choroid plexus   - 1 of 1 male (Grade 2) and 1 of 1 females (Grade 1) |  |
| Lymphohistiocytic inflammation extended into the vasculature of the brain parenchyma   - 1 of 3 males (5101; Grade 1) and 0 of 3 females |  |  |
| **Group 4** | |  |
| **Scheduled necropsy on day 30** | **Scheduled necropsy on day 72** |  |
| Lymphoplasmacytic inflammation of the choroid plexus   - 1 of 3 males (Grade 1) and 2 of 3 females (Grade 1) | Lymphoplasmacytic inflammation of the choroid plexus   - 1 of 2 males (Grade 1) and 1 of 2 females (Grade 1) |  |
| **Group 2, 3** | |  |
| - | - | - no-observed-adverse-effect level; area under the curve[0-168h]) 474,000 µg•h/mL |

Table S6. Quantification of LAG-3 expression by real-time PCR.

| **Sample** | **PPIA** | **LAG-3 CT** | **Mean PCT CT normalized to PPIA (Δ CT)** |
| --- | --- | --- | --- |
| Pituitary, in-house | 19.09 | 33.06 | 13.96 |
| Pituitary, commercial | 19.72 | 34.42 | 14.71 |
| MNCs day 0 | 17.26 | 31.98 | 14.72 |
| T cells | 22.06 | 31.10 | 9.05 |
| NTC | NS | NS |  |

The mean PCR cycle time (CT) of each sample was normalized to *PPIA* (cyclophilin A, housekeeping gene). The relative abundance was then determined by normalization to the CT for bone marrow mononuclear cells (MNCs). The results showed that, relative to *PPIA*, the abundance of *LAG-3* mRNA in the examined tissue RNA samples was lower in pituitary compared to CD8+ T cells but approximately equal to that found in the MNCs.

### Supplementary Figures

Figure S1.

Binding of relatlimab to LAG-3 recombinant and cell-surface LAG-3. **A**, Binding of relatlimab to human LAG-3 by ELISA (left panel) and flow cytometry (right panel). **B**, Surface plasmon resonance sensogram depicting binding of monomeric relatlimab Fab under acidic pH conditions (pH 6). MFI, mean fluorescence intensity; OD, optical density; RU, response units.

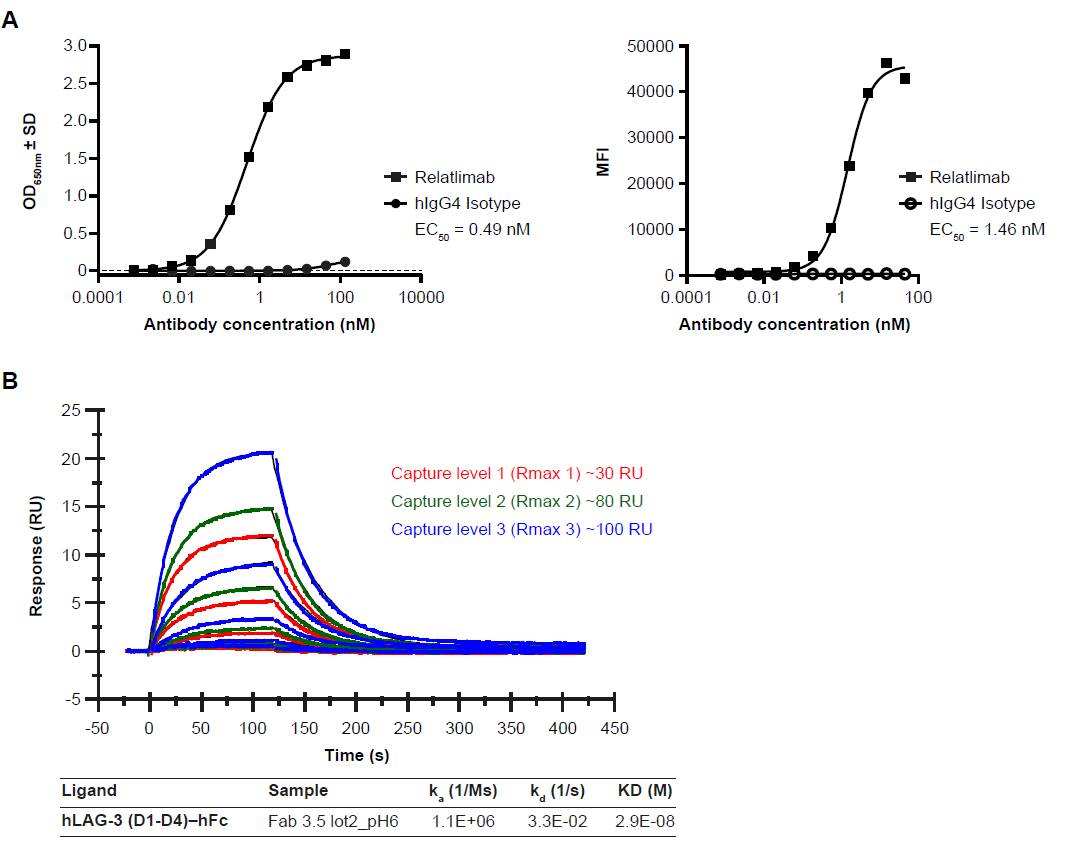

Figure S2

Interaction of LAG-3 and MHC II, comparing binding to **A**, wild-type and **B**, mutant MHC II-low variant RJ225 of the Raji B cell lymphoid cell line. **C**, Flow cytometric characterization of MHC II expression in each cell line. FL2-GMFI, geometric mean fluorescence intensity; mLAG-3, mouse LAG-3.

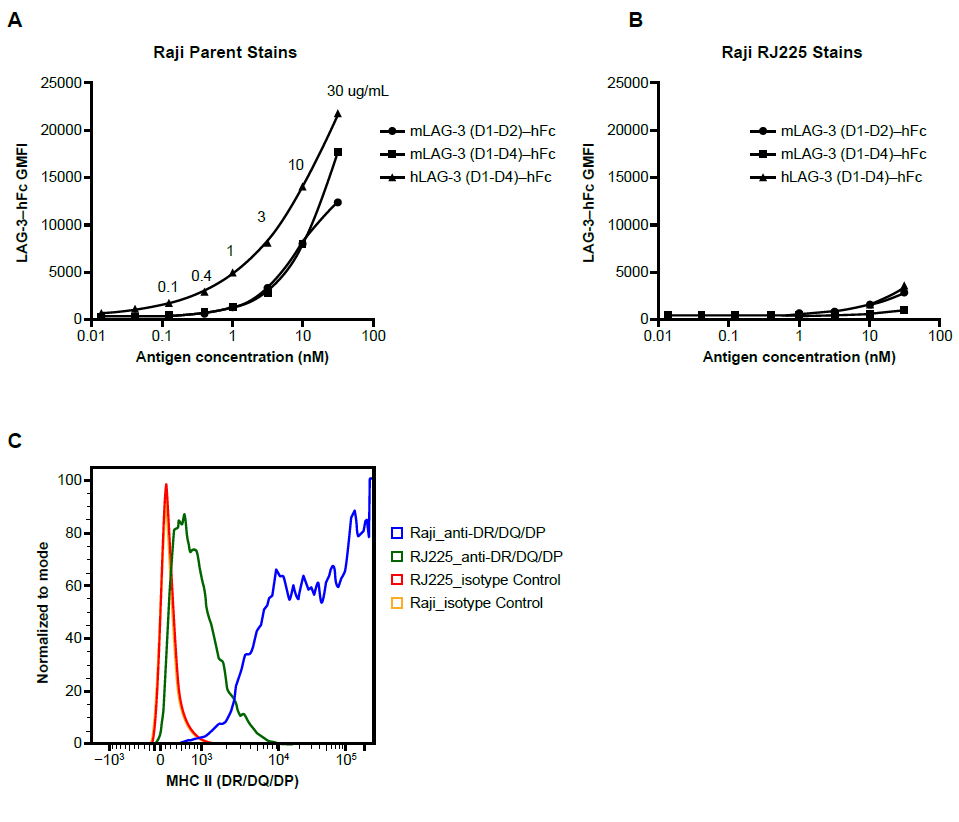

Figure S3.

Octet^®^ biolayer interferometry measurements. **A**, Blockade of LAG-3 binding to HLA-DR. **B**, LAG-3 binding to HLA-DR in the presence of relatlimab and D3–D4-specific LAG-3 antibodies. **C**, Determination of human LAG-3 domain specificity of LAG-3 monoclonal antibodies. Ab, antibody.

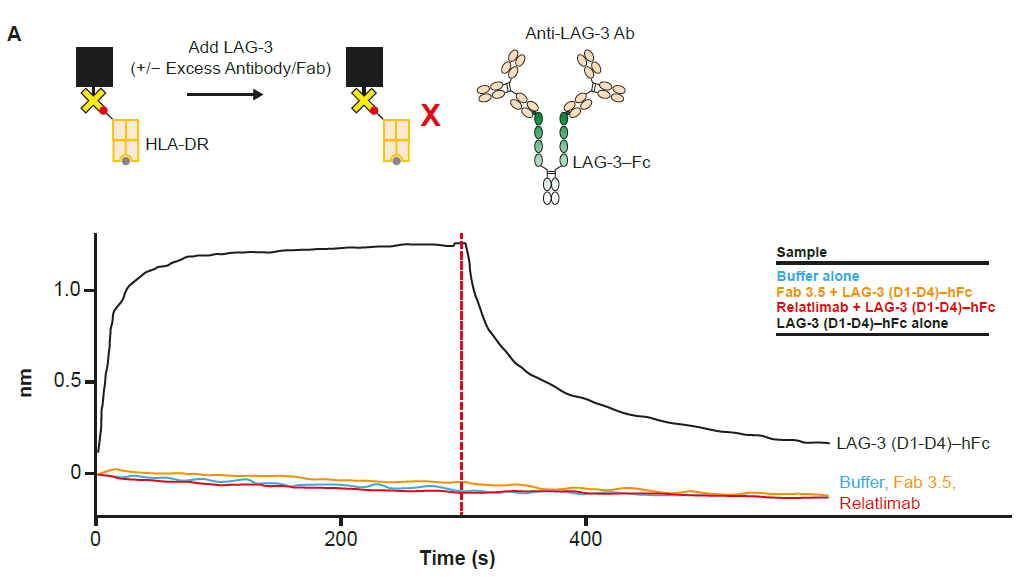

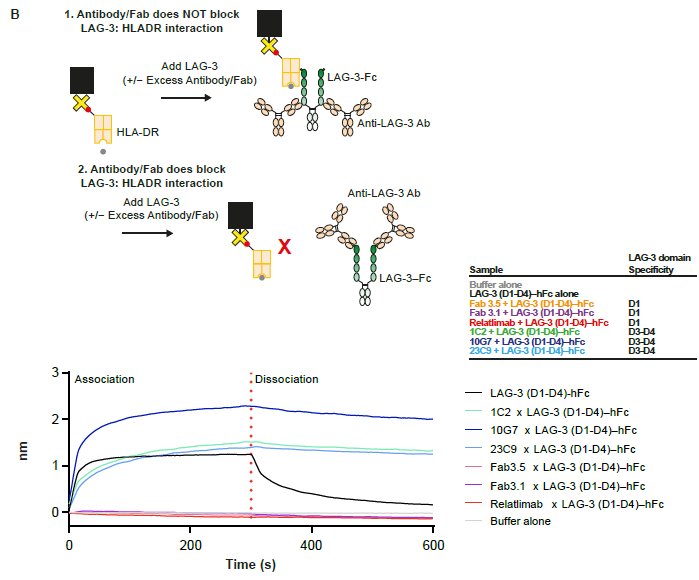

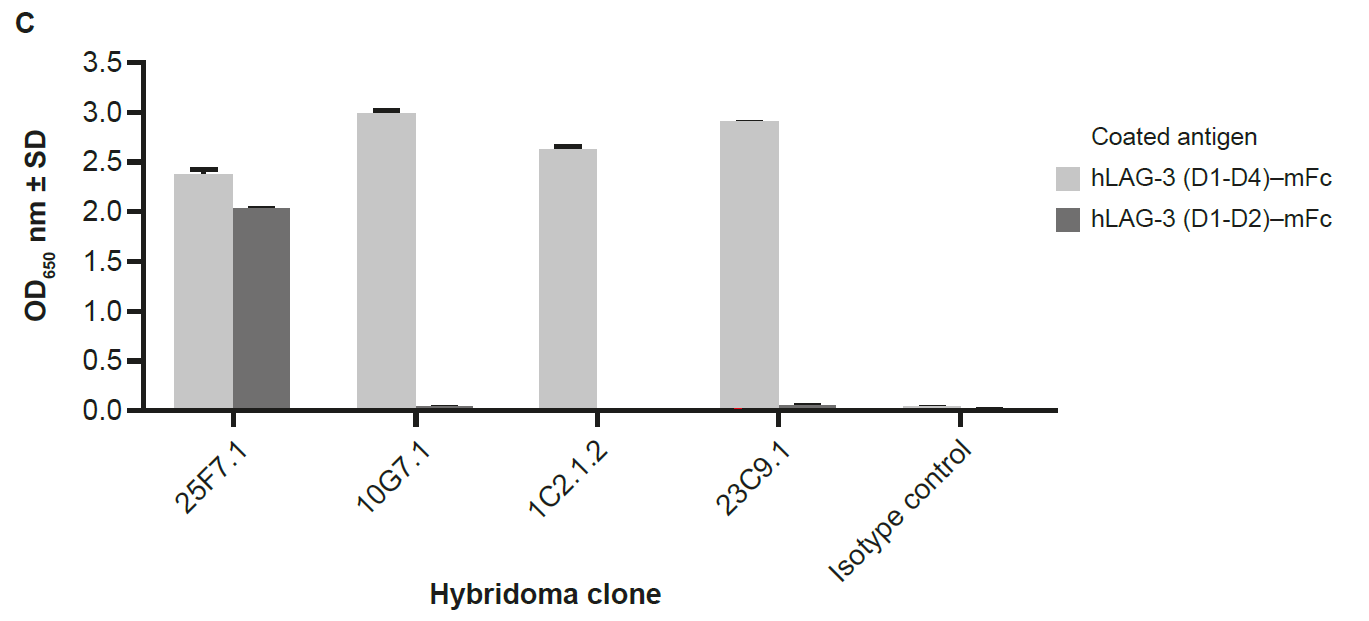

Figure S4.

Limited inhibition of T-cell responses by soluble hexameric FGL1-RGY in the 3A9 T-cell hybridoma model. FGL1-RGY, fibrinogen-like protein-1 fused with hexamer-forming Fc (E345R/E430G/S440Y); mIL-2, mouse interleukin 2.

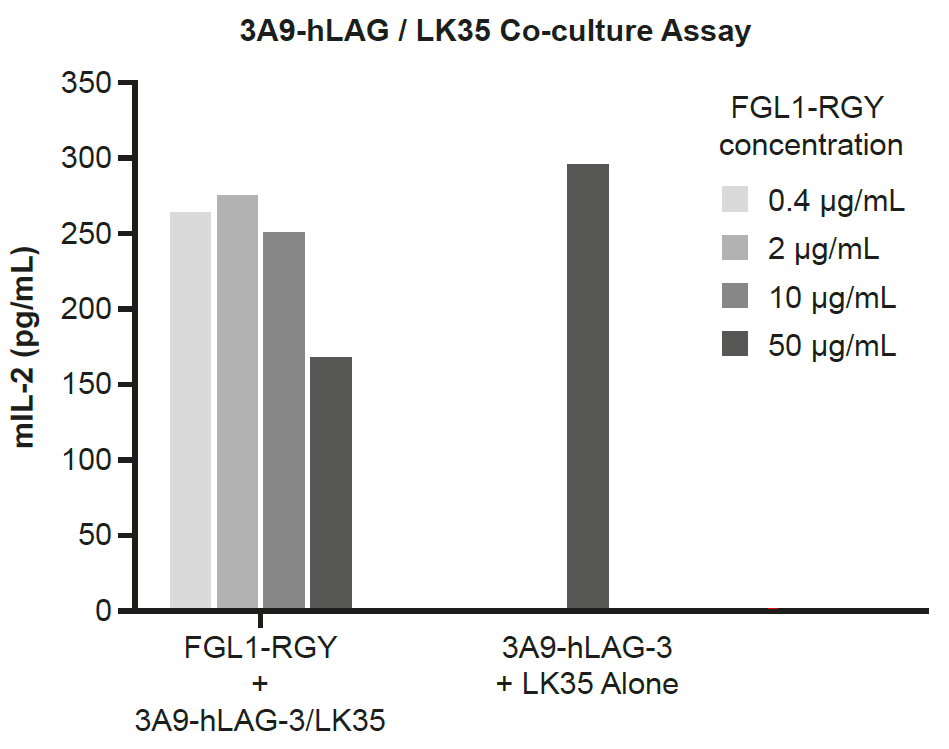

Figure S5.

Components of the LAG-3 functional cell-based bioassay exploring FGL1 engagement of LAG-3. **A**, Co-culture of 3A9-hLAG-3 T cells and LK35 APC engineered for co-expression of both endogenous MHC II and ectopic FGL1_MT._ **B**, Flow cytometric analysis of FGL1 and MHC II expression on the LK35 parent and LK35-FGL1_MT_ APC lines. APC, antigen-presenting cell; MT, membrane tethered; mCD4, murine CD4.

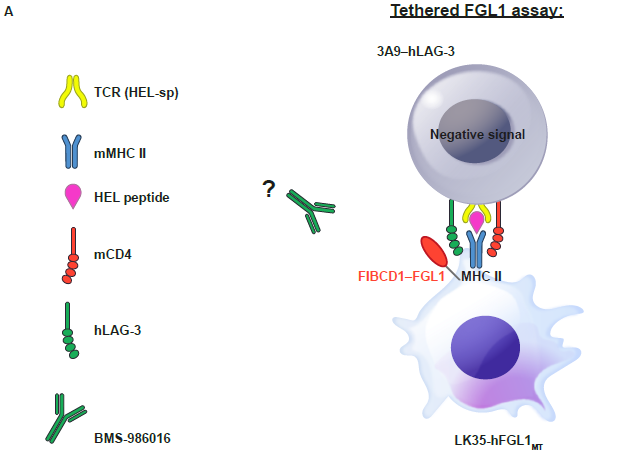

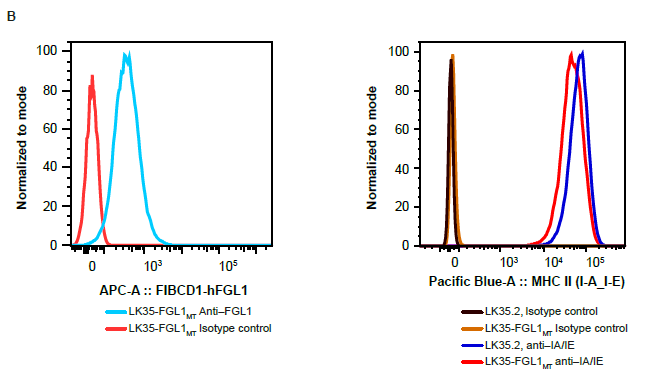

Figure S6.

Evaluation of relatlimab-mediated ADCC on activated primary human T cells from two donors. **A**, Donor A. **B**, Donor B.

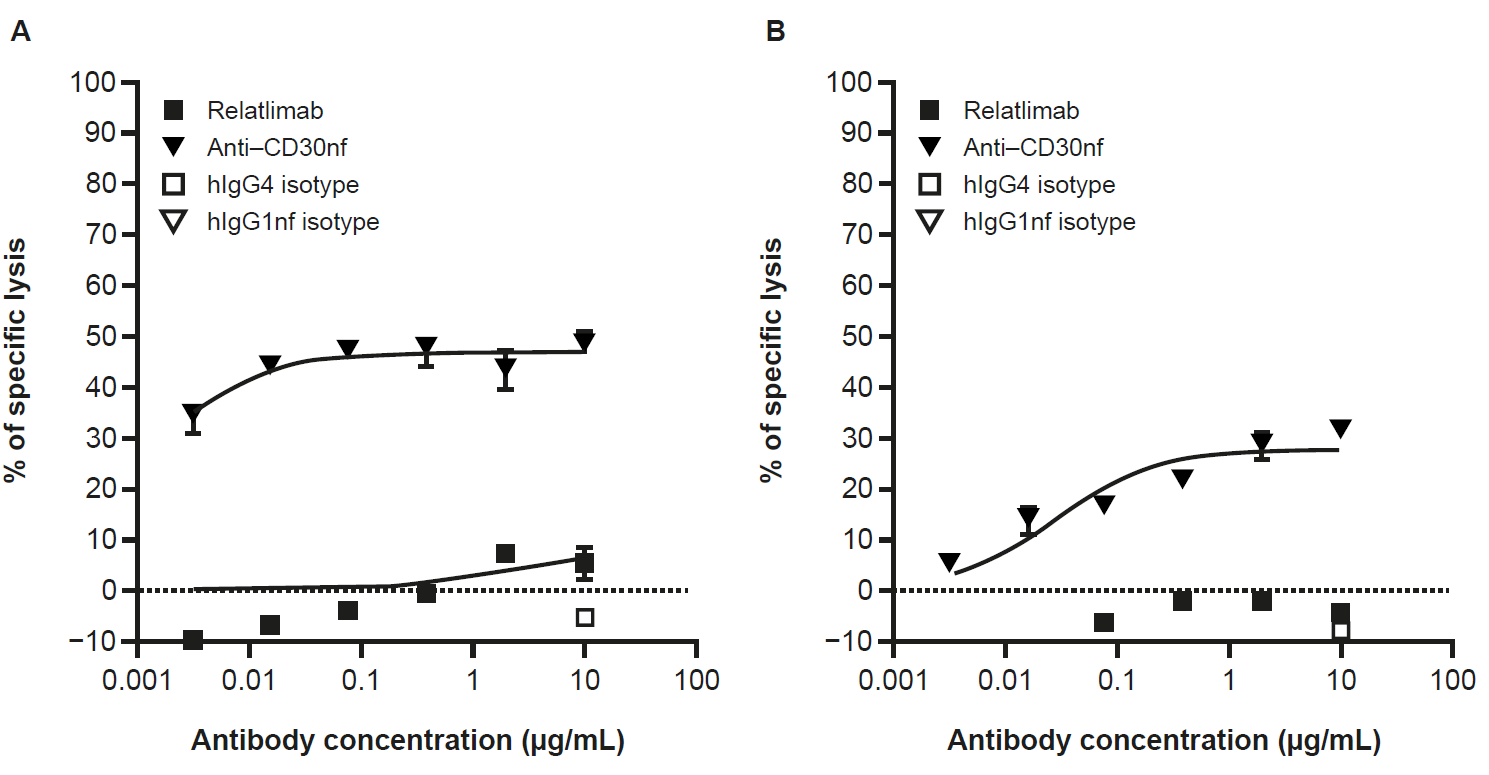

Figure S7.

Antitumor activity of anti–LAG-3 antibody alone and in combination with anti–PD-L1 in an MC38 colon adenocarcinoma mouse model in C57BL/6 mice. **A**, Mean tumor volumes across treatment groups. **B**, Individual mouse tumor growth curves.

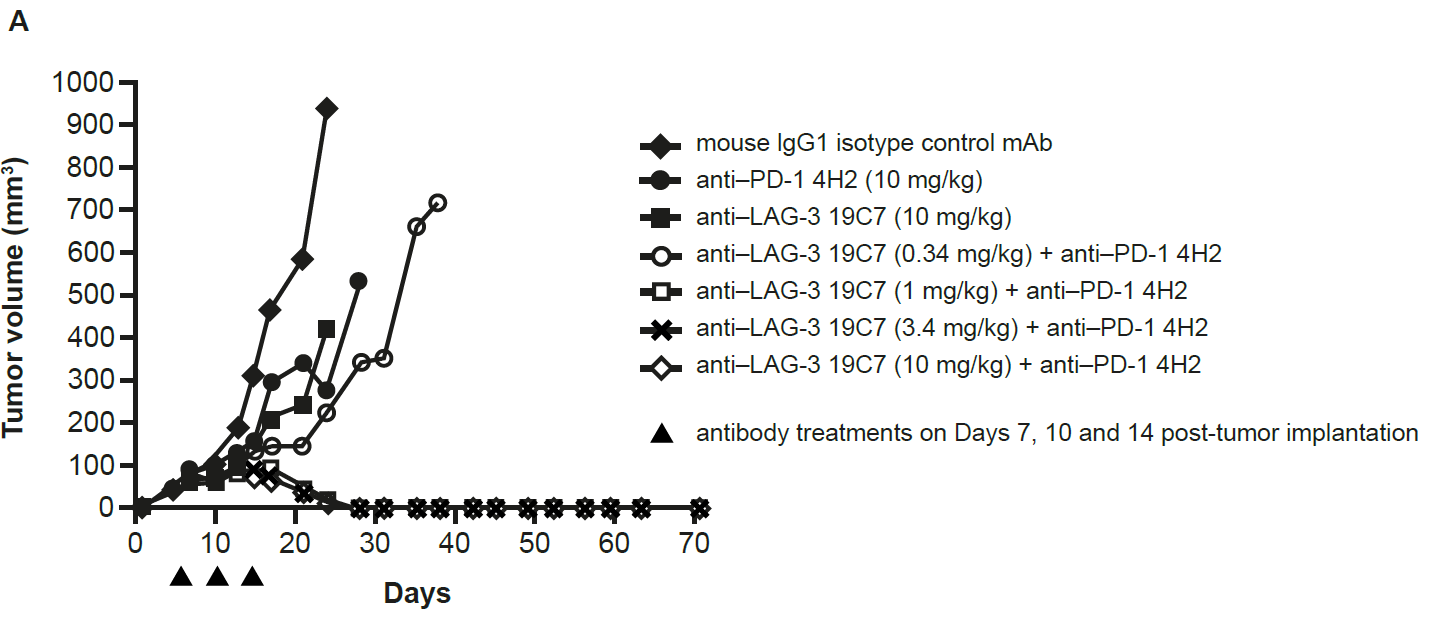

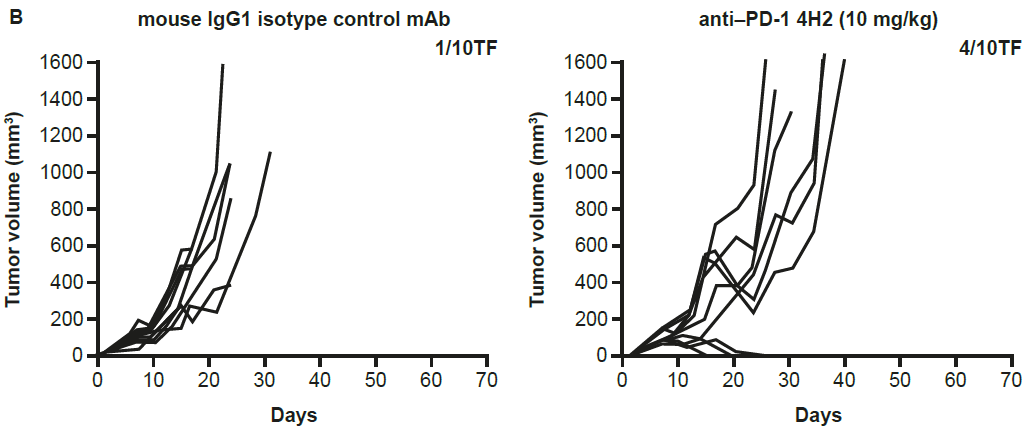

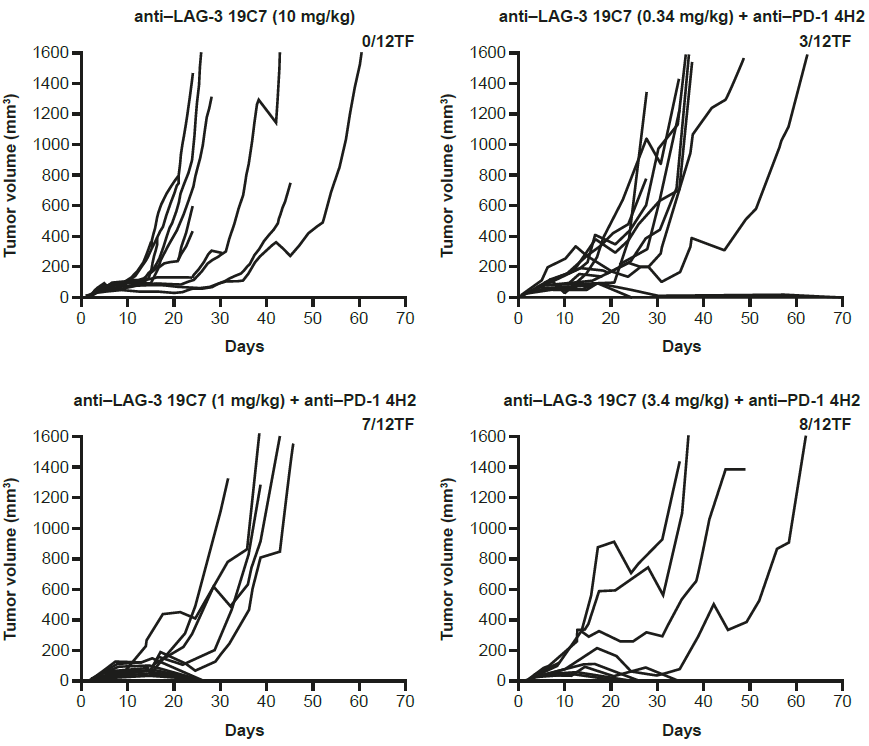

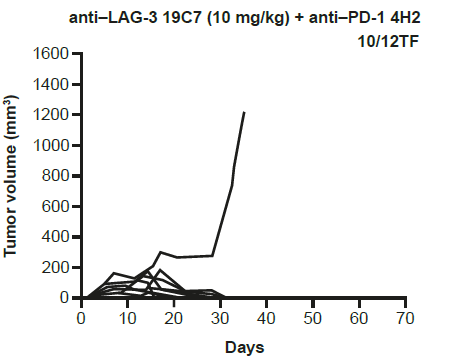

Figure S8.

Antitumor activity of anti–LAG-3 antibody alone and in combination with anti–PD-1 in an SA1N fibrosarcoma model in A/J mice. **A**, Mean tumor volumes across treatment groups. **B**, Individual mouse tumor growth curves.

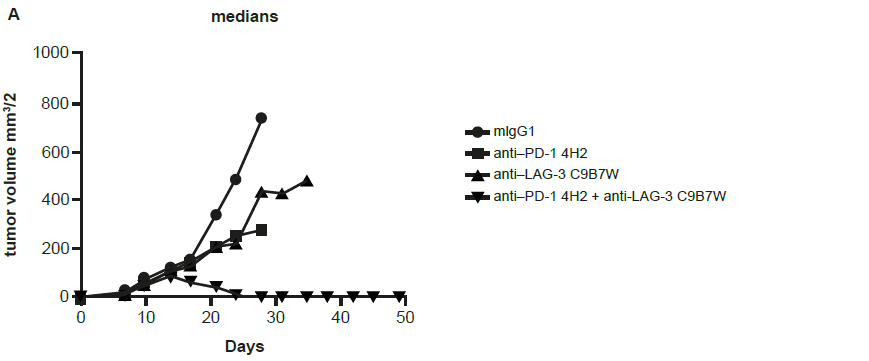

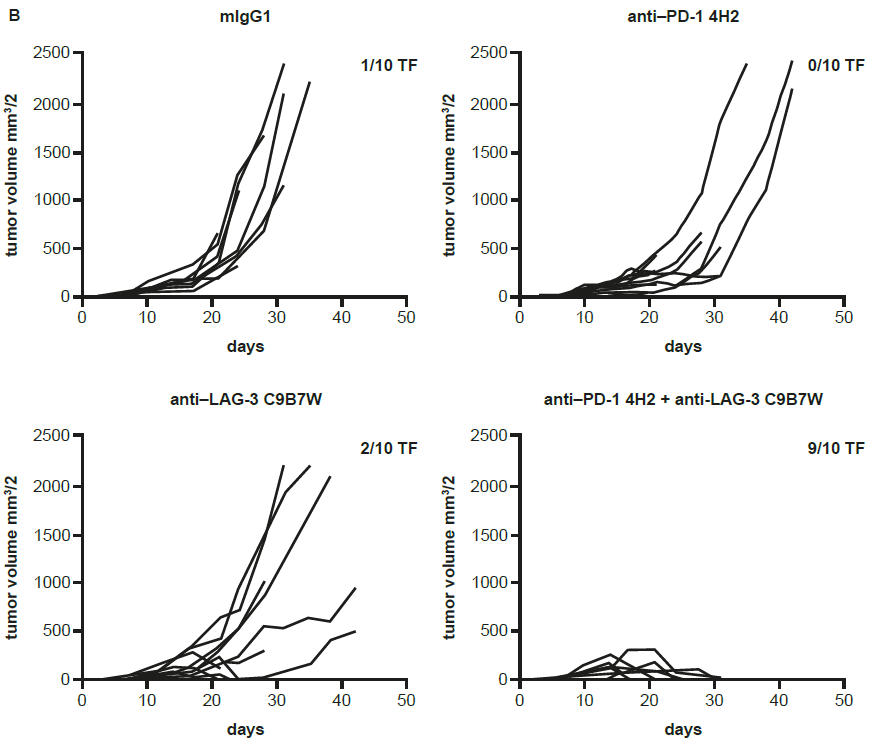

Figure S9.

**A**, Overlapping peptides spanning the LAG-3 insertion loop. **B**, Relatlimab epitope and its conservation across species. **C**, Design rationale for hLAG-3 loop variants.

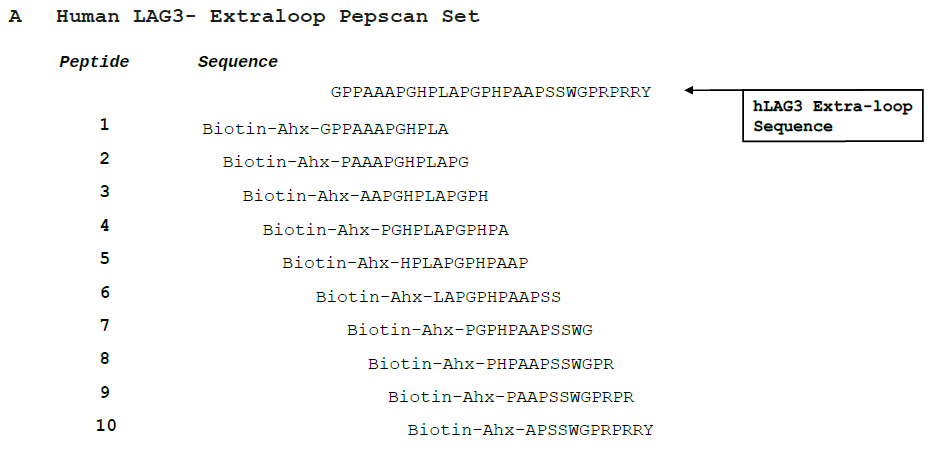

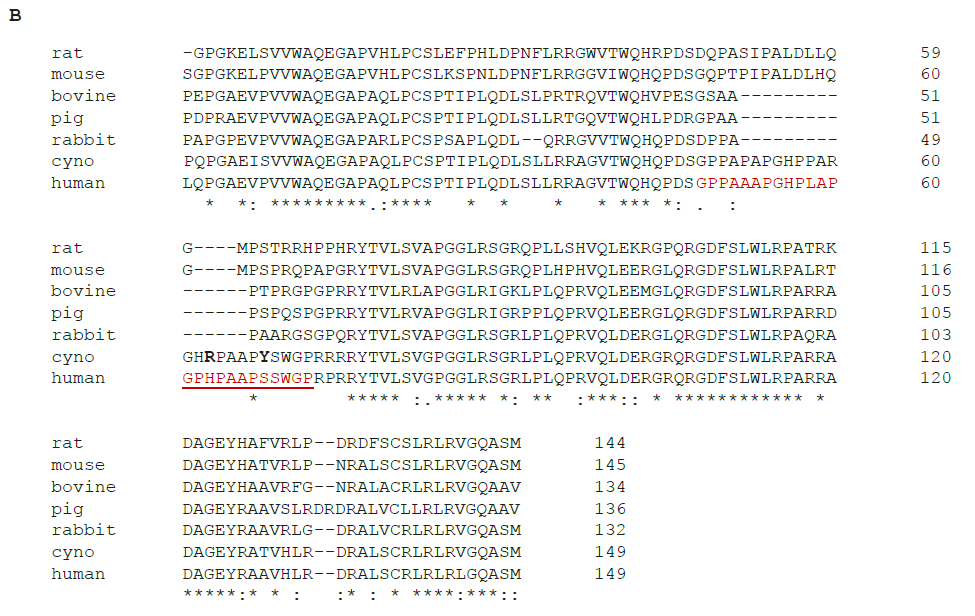

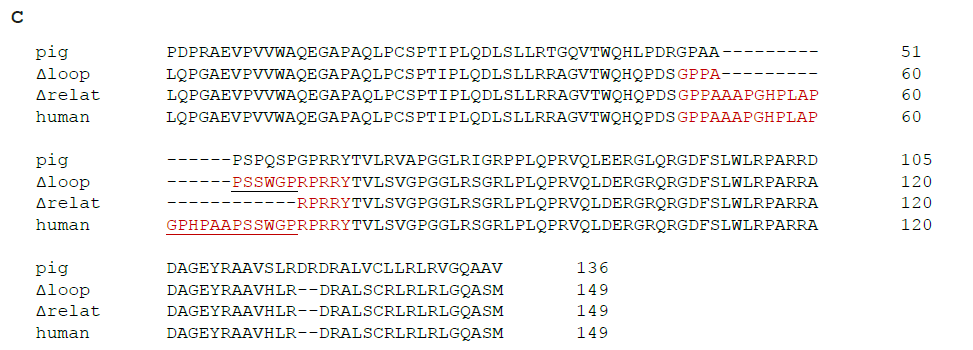

Figure S10.

Human LAG-3 fusion protein blocks the binding of HuMAb anti–LAG-3. Pre-incubation of LAG-3.1-G4P-FITC (10 µg/mL for immunofluorescent and 5 µg/mL for immunoperoxidase method) with five-fold excess of human LAG-3 fusion protein for 2 hours before applying to pituitary sections. Top and middle panels represent immunofluorescent staining and bottom panels represent immunoperoxidase staining.

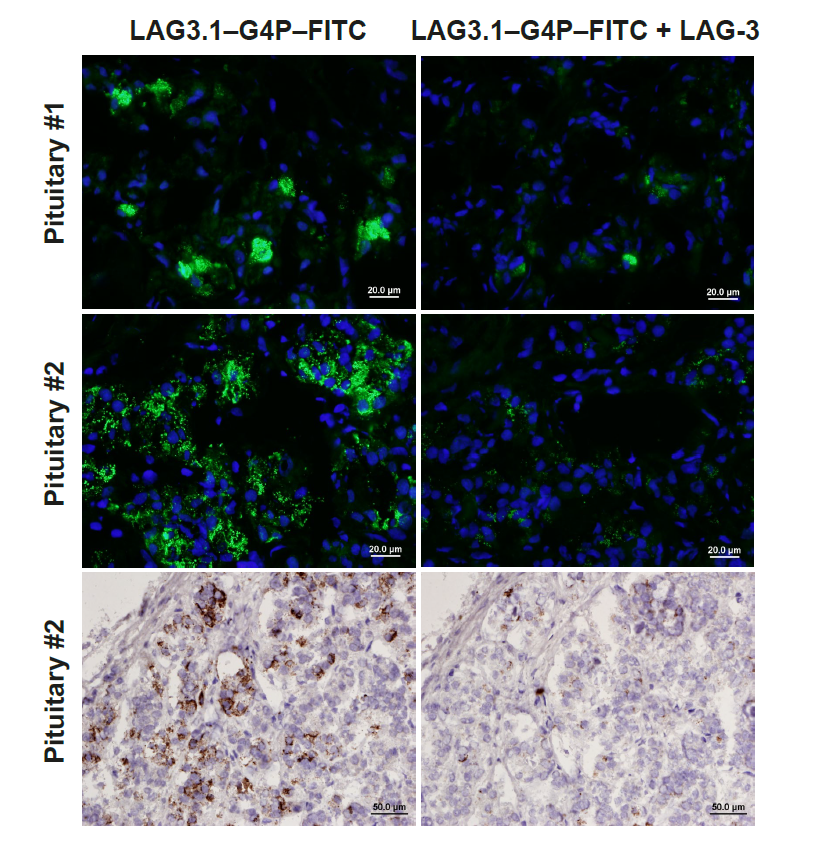

Figure S11.

**A**, LAG-3 RT-PCR products detected by agarose gel electrophoresis. **B**, PCR primer map for human LAG-3 gene and respective transcript.

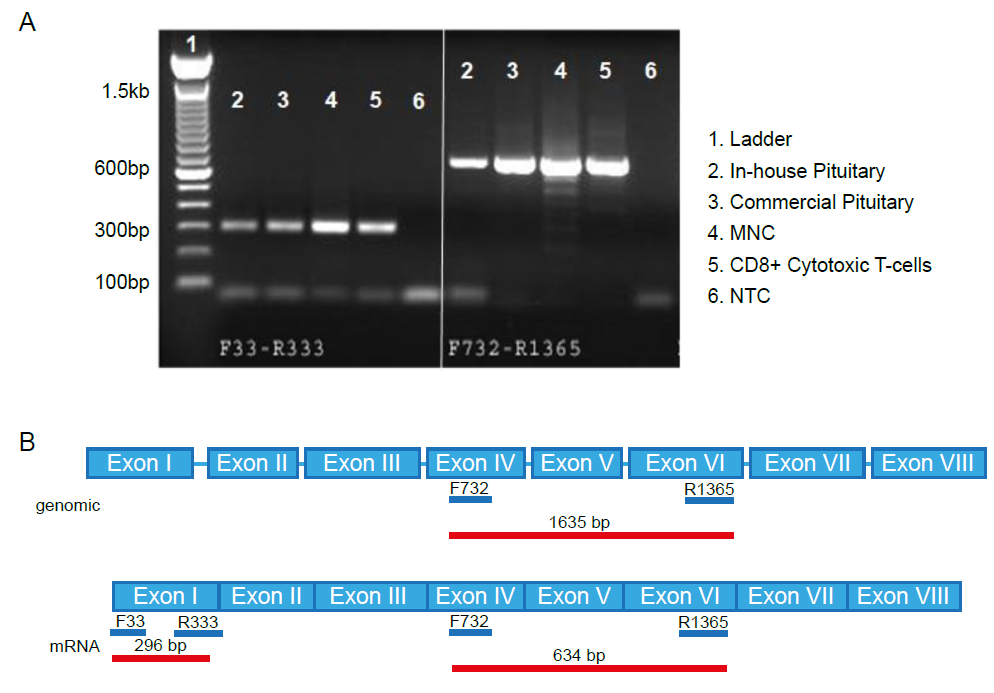

The presence of LAG-3 expression in human pituitary tissue was confirmed using PCR. The primer pair generated a PCR product length consistent with amplification of hLAG-3 mRNA (634 base pairs) but did not generate a band of 1635 base pairs, demonstrating absence of contaminating genomic DNA in sample preparation. In all cases, no products were detected in the non-template control.

**Figure S12.**

Histopathology from a single male monkey treated with relatlimab 100 mg/kg + nivolumab 50 mg/kg euthanized in moribund condition on day 29. Images show slight to moderate lymphoplasmacytic inflammation of the choroid plexus (left) and mixed-cell inflammation of the epididymis (right).

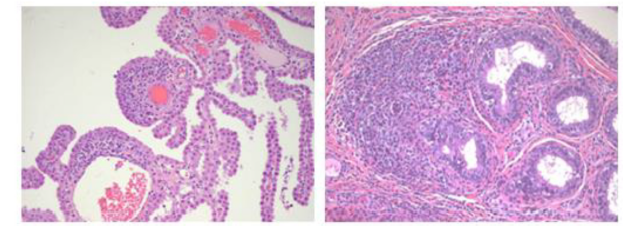

Figure S13.

Splenic T-lymphocyte phenotyping analysis of 4-week monkey toxicity study animals. The percentage of TNFα^+^, INFγ^+^, and CD69^+^ CD4^+^ CD8^−^ T cells from 1–5 **A,** male and **B,** female animals per group are shown for each indicated timepoint (mean ± SEM). SEM, standard error of the mean.

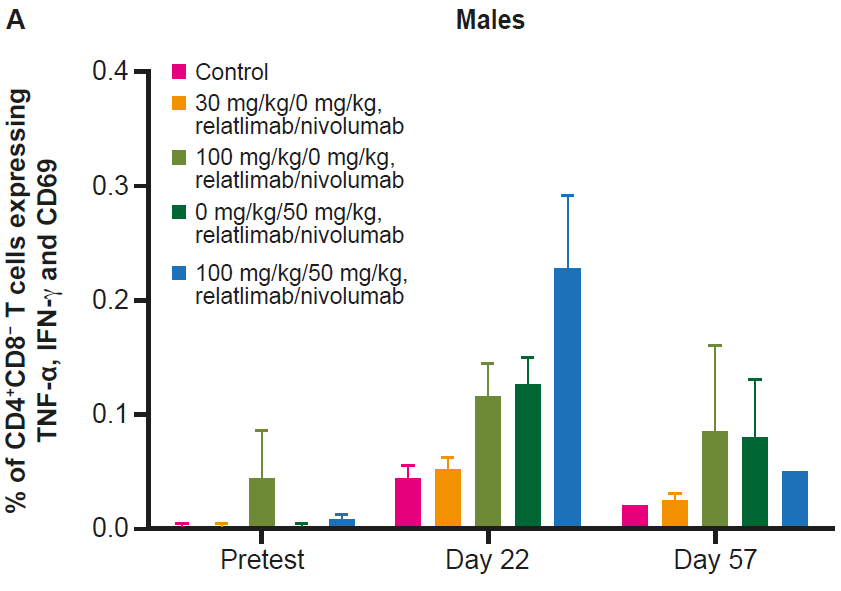

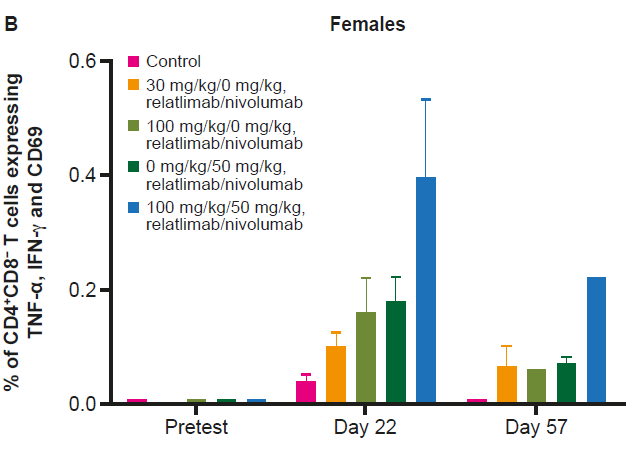

Figure S14.

Splenic T-lymphocyte phenotyping analysis of 4-week monkey toxicity study animals. The percentage of CD4^+^ T-regs from 1–5 **A**, male and **B**, female animals per group are shown for each indicated timepoint (mean ± SEM). T-regs, T regulatory cells.

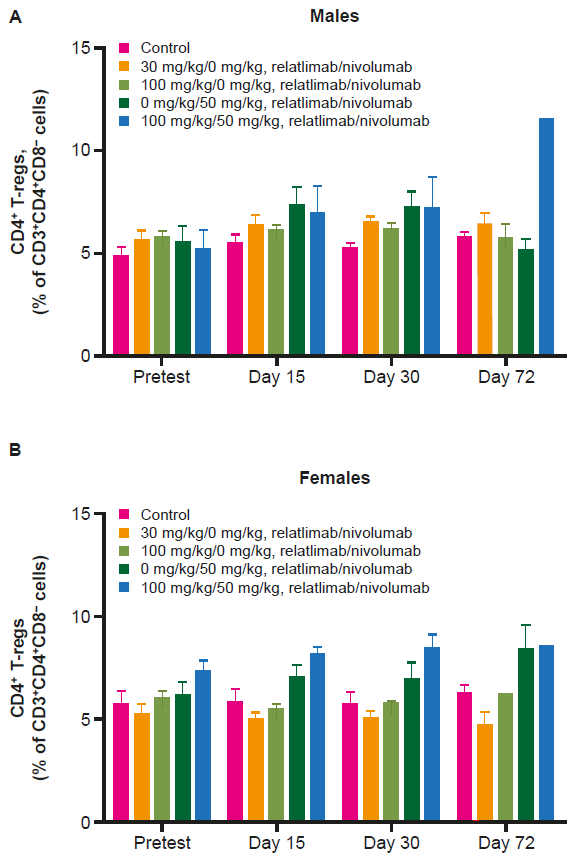
